# Conditional activation of NK cell function using chemically synthetic constrained bicyclic peptides directed against NKp46 and tumor-expressed antigens

**DOI:** 10.1101/2024.10.30.621052

**Authors:** Fay J. Dufort, Christopher J. Leitheiser, Alexandra Rezvaya, Tucker R. Ezell, Kathleen Q.W. Ho, Gustavo A. Bezerra, Ben J. F. Blakeman, Sandra Uhlenbroich, William H. Zammit, Lukas Stanczuk, Peter N. Brown, Gemma E. Mudd, Kevin McDonnell, Nicholas Keen, Philip E. Brandish

**Affiliations:** Bicycle Therapeutics, Inc. 35 Cambridgepark Drive Suite 350 Cambridge, MA 02140, USA; BicycleTx Limited, Portway Building, Granta Park, Cambridge CB21 6GS, UK

**Author notes:** Correspondence: PEB. Conflict of interest disclosure statement: All authors were full-time employees of Bicycle Therapeutics at the time that the work was conducted, and some own stock or stock options in Bicycle Therapeutics.

## Abstract

Natural killer (NK) cells have the unique potential to recognize and kill tumor cells independently of MHC-I presentation of antigens, as well as to secrete cytokines that engage adaptive anti-tumor immunity and the function of cytolytic T cells. We have discovered and characterized chemically synthetic, constrained bicyclic peptides that bind with high affinity and specificity to NKp46, an activating receptor expressed selectively on NK cells in the tumor microenvironment. Chemical coupling to other bicyclic peptides specific for the tumor antigens EphA2 or MT-1 created NKp46 agonists whose function was completely conditional on binding to the tumor antigen. These chemical conjugates effectively convert the tumor antigen into a “kill me” signal for NK cells. Not only did these newly created tumor-immune cell agonists (TICAs) direct potent and efficient killing of human tumor cells by primary human NK cells in vitro, but they also caused secretion of the pro-inflammatory cytokines TNFα and IFNγ. Importantly, the TICAs directed production of FLT3 ligand, an essential mitogen for conventional dendritic cells which are central to the development of anti-tumor immunity in cancer. We illustrate the TICA-directed interaction of NK cells with tumor cells using confocal microscopy and we show that TICAs enable sustained function over multiple rounds of killing. These novel tools are well positioned to harness the potential of NK cells in the treatment of cancer.

## INTRODUCTION

Natural killer (NK) cells are so called because of an innate ability to surveille, recognize and kill stressed, infected or transformed (cancer) cells without prior exposure to a particular antigen (1–4). This stands in contrast to cytolytic T cells which require priming and presentation on the target cell surface of a specific antigen in the context of MHC-I. NK cells detect loss of or aberrant MHC-I expression and other cellular stress markers via invariant activating and inhibitory surface receptors whose signals integrate intracellularly to drive cytolytic function(1). It is self-evident that cancers that present clinically have escaped this immune surveillance function as well as adaptive immune control. NK cells are nevertheless attractive as point of therapeutic intervention in cancer because they sit at the interface of innate and adaptive immunity, killing tumor cells without prior exposure to specific antigens, making cancer antigens available to macrophages and dendritic cells, and producing cytokines that alert and activate adaptive immune cells(5–8). Numerous studies have documented the presence of NK cell infiltrates in a range of major underserved tumor types(9–13), including very recent publications that have dissected the phenotypes of those cells in considerable detail(14–16). We and others hypothesized that localized and specific pharmacological activation of NK cells in situ could trigger a fresh wave of antitumor immunity that could offer meaningful clinical benefit to cancer patients, especially in combination with existing therapeutics. Likewise, there is considerable interest in employing NK cells themselves as therapeutics on account of their allogenic nature and potentially better safety profile compared to T cell therapy(17–19).

As noted above, NK cells use an array of surface receptors to interrogate the status of potential target cells. We selected the activating receptor NKp46(8,20,21) for the construction of a tumor-targeted immune cell agonist (TICA) using chemically synthetic constrained bicyclic peptides to bind to and conditionally activate NK cells in the presence of specific tumor antigens (NK-TICA). NKp46 is expressed selectively and universally on both resting and activated NK cells(22), so much so that it is used a marker for tumor-infiltrating NK cells, and expression seems to preserved in the tumor unlike CD16 and NKG2D(12). This lack of down-regulation suggested to us that NKp46 is generally not activated in the tumor setting and is therefore available to drive NK cell function. Others have described bi- or trispecific cell engagers targeting NKp46 both for solid tumors and for hematologic cancers(12,23–26), with some progressing into the clinic and at least one affording clinical responses(27,28).

The Bicycle® platform first described by Heinis et al.,(29) offers potential advantages in the setting of immune cell engagers. The concept is exemplified by the discovery and development of a Nectin-4-targeted conditionally active CD137 agonist(30,31), including small size for rapid penetration into tissues(32) (Eder, 2019), completely conditional function, and a short controllable half-life in the body(31). We also surmised that the flexible formatting and small size of TICAs would avoid any reliance on co-activation of CD16, minimizing the risk of NK cell fratricide, and enable optimal dosing to avoid the potential for NK cell exhaustion. In this report, we describe the initial in vitro proof of concept for reprogramming of tumor antigens as a “kill me” signal for NK cells via the activating receptor, NKp46.

## METHODS

### Cell isolation

NK cells were isolated from whole blood (StemExpress catalog #PBEDT450F) or leukopaks (AllCells, catalog #LP-CR-CD56+-NS-10M). Blood is diluted 1:1 with PBS and PBMC isolated by density centrifugation in Sep-Mate 50 IVD Tubes (StemCell catalog #85450) over Ficoll-Paque™ PLUS (Fisher Scientific 4 catalog #5-001-749). NK cells were then following EasySep™ Human NK Cell Isolation Kit (StemCell catalog #17955) per manufacturer’s instructions. NK cells were utilized immediately in an in vitro assay or stored in cell freezing media (CryoStor catalog #CS10) at ∼10 million/mL for short term storage in liquid nitrogen.

### NK-mediated tumor cell killing

The tumor cell lines HT1080-luc cells (ATCC, catalog #CCL-121-LUC2 or Incucyte® catalog #4486) or A431-luc (ATCC, catalog #CRL-1555-LUC2) were grown and maintained according to manufacturer’s recommendation. At the time of tumor killing assay, 20000 NK cells were co-cultured with 4000 HT1080-luc cells or 2000 A431-luc cells in a 96 well flat bottom optical plate. Culture media contained RPMI-1640 (Gibco™ catalog #11875-093; with L-glutamine) with 10 % heat-inactivated fetal bovine serum (FBS; Corning® catalog #35-011-CV), 10 mM HEPES (Gibco™ catalog #15-630-080), and 1% Penicillin Streptomycin (Corning® catalog #30-002-CI) with a final concentration of 50 IU/mL of IL-2 (Miltenyi Biotech catalog #130-097-743). After 24 hr incubation at 37 °C, 5% CO_2_, luciferase levels were measured by Bright-Glo™ Luciferase Assay System (Promega Corporation catalog #E2620) with a luminometer (CLARIOstar®, BMG Labtech). Data was analyzed with GraphPad Prism® with a 4-parameter kinetic fit.

### Cytokine secretion assay

NK cells were co-cultured with 40,000 HT1080-luc tumor cells at a 5:1 effector to target ratio for 4 hr or 48 hr 37 °C, 5% CO2, in RPMI-1640 (Gibco™ catalog #11875-093; with L-glutamine) with 10% heat-inactivated fetal bovine serum (FBS; Corning® catalog #35-011-CV), 10 mM HEPES (Gibco™ catalog #15-630-080), and 1 % Penicillin Streptomycin (Corning® catalog #30-002-CI) with a final concentration of 50 IU/mL of IL-2 (Miltenyi Biotec catalog #130-097-743). Cell-free supernatants were collected, cytokines measured by MesoScale Discovery (MSD) platform (interferon-gamma and tumor necrosis factor-alpha, catalog #K15067M-2) or by ELISA (Human Flt-3 Ligand/FLT3LG Quantikine ELISA Kit; RnD systems catalog #DFK00) according to manufacturer’s instructions. Data was analyzed with GraphPad Prism® with 4-parameter kinetic fit.

### Flow cytometry

PBMCs and NK cells were isolated and plated in a 96-well V-bottom polypropylene plate (Greiner Bio-One catalog #651201) in RPMI-1640 (Gibco™ catalog #11875-093; with L-glutamine) with 10% heat-inactivated fetal bovine serum (FBS; Corning® catalog #35-011-CV), 10 mM HEPES (Gibco™ catalog #15-630-080), and 1% Penicillin Streptomycin (Corning® catalog #30-002-CI). NK cells and/or PBMCs were then stained with viability dye (eBiosciences catalog #65-0865-18) followed by fluorochrome conjugated antibodies for 30 minutes, followed by Bicycle peptides fluorescently conjugated with AF647 (Click Chemistry Tools, Beijing, China, or Lumiprobe Corporation, USA catalog #66820) for 1 hr. Samples were then acquired on a BD Symphony™ A3 flow cytometer and analyzed using FlowJo™ version 10 software. Relative receptor quantification was conducted on NK cells or tumor cells and determination of receptor numbers were calculated using the BD Quantibrite™ PE Phycoerythrin Fluorescence Quantitation Kit catalog #340495 and manufacturers’ protocol. Data was analyzed with GraphPad Prism® with 4-PL curve fit. Antibodies used in this manuscript can be found in Supplemental section 1.

### Peptide and conjugate synthesis

Bicyclic peptides were synthesized on Rink amide resin using standard Fmoc (9-fluorenylmethyloxycarbonyl) solid-phase peptide synthesis, either by manual coupling (for large scale) or using a Biotage® Syro II automated peptide synthesizer (for small scale), as previously described(29,30). Fluorescently labeled bicyclic peptides and NK-TICA molecules were also synthesized analogous to those described previously(30). Protocols can be found in Supplemental Section 2.

### Surface plasmon resonance

For analysis of binding affinities, a Biacore™ 8k+ instrument was used with a CM5 chip (Cytiva). SPR experiments were performed to determine k_a_ (M^-1^s^-1^), k_d_ (s^-1^), K_D_ (nM) values (where measurable) of compounds against immobilized NKp46 (Fc-tagged human NKp46 (ACRO Biosystems, catalog #NC1-H5257 and C365P1-202KF1-S3), EphA2(30), MT1(32). Sensorgrams and method can be found in supplemental information (Supplemental Section 1)

### Sequential killing assay

HT1080-Nuc Light (IncuCyte® Nuclight Green catalog #4486) were placed in co-culture with rested NK cells (overnight at 37 °C, 5 % CO2 in RPMI-1640 (Gibco™ catalog #11875-093; with L-glutamine) with 10 % heat-inactivated fetal bovine serum (FBS; Corning® catalog #35-011-CV) and 100 IU/mL of IL-2 (Miltenyi Biotec catalog #130-097-743), seeded in 96 well flat bottom plates (Greiner Bio-One catalog #655180) with media containing 50 IU/mL IL-2 at 10:1 effector to target cell ratio (20000: 2000). NK-TICA molecules (2 nM) were added to the culture at initiation. Additions of HT1080-Nuclight target cells were daily in RPMI-1640 media containing 50 IU/mL IL-2 for a daily final volume of 100 µL/well. Cultures were monitored in Incucyte® S3 live cell imaging system, 1-4 images per well, every 3 hr. Images were processed using Incucyte® software and analyzed with GraphPad Prism®.

### Imaging

Prior to imaging, HT1080 cells (ATCC; catalog #CCL-121-LUC2) were plated overnight in 8-well chambered glass slides (Nunc™ Lab-Tek™ II Chambered coverglass catalog #155409) in 250 µL/well of phenol free RPMI-1640 with 10 % FBS at 37 °C. NK cells were rested overnight at 37 °C in RPMI medium with 10 % FBS supplemented with 100 IU/ml recombinant human IL-2. The following day, NK cells were labeled with LysoView™ 594 (Biotium® catalog #70084T) and either Alexa Fluor® 647 anti-human NKp46 antibody (clone 9E2 Biolegend catalog #331910) or AF 647 (Lumiprobe Corporation, USA catalog #66820)-conjugated NK-TICA molecule. NK cells labeled with anti-human NKp46 antibody were incubated with unlabeled or non-binding (nb) NK-TICA molecules or left as untreated control. NK cells not incubated with anti-human NKp46 antibody were incubated with AF 647-labeld NK-TICA molecule. HT1080 cells were labeled with CellBrite®™ Steady 488 membrane dye (Biotium® catalog #30106-T) according to manufacturer’s instructions. NK cells were added to plated HT1080 cells in chambered slides with phenol free RPMI-1640 supplemented with 10 % FBS and 50 IU/ml recombinant human IL-2. Each well was imaged within 1 hr of co-culture with a spinning disk confocal microscope: Yokogawa CSU-X1 spinning disk confocal scan head, Nikon Ti-E inverted microscope stand with Perfect Focus System, motorized stage and DIC and phase optics, Prior® Lumen 200 metal halide light source, 4 laser lines in Andor ALC: 405/488/561/642 nm, 2 Andor iXon 897E back-illuminated EMCCD/CCD cameras (Oxford Instruments America Inc), using MetaMorph® acquisition software. Data was exported and analyzed using ImageJ software (Rasband, W.S., ImageJ, U. S. National Institutes of Health, Bethesda, Maryland, USA, https://imagej.net/ij/, 1997-2018).

### Protein expression and purification for crystallization

Briefly, pET22(b) vector harboring NKp46 construct 22-212 was used to transform chemically competent T7 Express cells (NEB, US) following protocols provided by the manufacturer. Transformants were plated onto 2YT agar plates supplemented with 100 µg/mL ampicillin and plates incubated at 37 °C. The following day, a single colony was picked and used to inoculate 100 mL 2YT media supplemented with 100 µg/mL ampicillin, cultures grown at 37 °C, shaking 200 RPM. The following day, 10 mL of culture was used to inoculate 1 L 2YT plus 100 µg/mL ampicillin media (total 8 L grown), cultures grown to an OD_600_ = 1.0, 37 °C, shaking 200 RPM and cultures induced with 1 mM IPTG. Cultures were left to grow for an additional 4 hr, harvested via centrifugation at 4000 RPM for 15 min, ambient temperature, and the cell paste washed once with PBS (100 mL/L culture paste) and suspension centrifuged again at 4000 RPM, 15 min, 4 °C. Cell paste was flash frozen in liquid nitrogen and stored at −80 °C.

For protein purification, refolding was performed. Cell paste was defrosted on ice, resuspended in 50 mL of Wash Buffer (50 mM Tris-HCl, 300 mM NaCl, pH 8.0) supplemented with BaseMuncher® Endonuclease, following recommendation from the manufacturer (Abcam, UK) and Sigma-Fast™ EDTA Free protease inhibitor cocktail (Sigma-Aldrich, UK). Cells were lysed using French press, 30 kPsi, passing once at 4 °C. Cellular debris and insoluble protein fractions were isolated via centrifugation at 50000 x g at 4 °C, 30 min. The insoluble fraction was then resuspended in Denaturation Buffer (50 mM Tris-HCl, 8 M urea, pH 8.0) at 10 mL/g of insoluble fraction.

Protein samples were then further diluted with Denaturation Buffer to achieve a protein concentration of 1-5 mg/mL for ensuring higher protein yields in rapid refolding step. Cell debris was then removed via centrifugation, 50000 x g at 20 °C, 30 min. The unfolded protein sample was rapidly diluted into Refolding Buffer (100 mM Tris-HCl, 0.4 M arginine, 5 mM cysteamine, 1 mM cystamine) to achieve final protein concentration of 0.05 - 0.1 mg/mL. Refolding was performed at 4 °C, solution rapidly stirred, and protein added using a syringe pump operated at a flow rate of 0.2 mL/min. Some precipitation was evident, and the refolding was allowed to take place for an additional 16 hr, 4 °C. Sample was buffer exchanged into Buffer A (20 mM Tris-HCl, 150 mM NaCl, pH 8.0) via diafiltration (Cytiva, UK). Protein samples were concentrated, aggregates removed by passing through 0.4 µm PES filter (Sartorius) and sample incubated with 2 NiNTA resin (Qiagen, USA). Protein samples were allowed to bind for 4 hr, resin harvested via centrifugation (800 RPM, 5 min, 4 °C) and used to pack Diba OmniFit® column (Cole-Parmer, UK) which was then attached to ÄKTA go™ FPLC (Cytiva Life Sciences, UK) equilibrated with Buffer A and Buffer B (20 mM Tris-HCl, 150 mM NaCl, 500 mM imidazole, pH 8.0). The column was washed with Buffer A, 2 % Buffer B and the protein eluted by applying 100 % Buffer B. Eluent was then concentrated and further purified by size exclusion chromatography (SEC), Superdex75 16/600 column (Cytiva, UK), equilibrated with Buffer A. A single major peak of > 95 % pure NKp46 was obtained, protein concentrated to 10 mg/mL, flash frozen in liquid nitrogen and stored −80 °C for future studies.

### Protein Crystallization and Structural Determination

Purified human NKp46 protein at 10 mg/mL in storage buffer (20 mM Tris-HCl, 150 mM, pH 8.0) was mixed with bicyclic peptides (50 to 100 mM in 99.9% pure, anhydrous DMSO, Sigma-Aldrich catalog number 276855) at a molar ratio of 1:1 or 1:2 and incubated for 120 min, on ice.

Initial crystal screening of the co-complex was performed by the vapor diffusion, hanging-drop method using commercially available sparse matrix screens including PACT Premier, Ligand Friendly Screen (LFS), Structure Screen 1+2, SG1 Screen and Morpheus (Molecular Dimensions, UK). A mosquito® liquid handler (SPT Labtech) dispensed either 100 nL: 100 nL or 200 nL: 100 nL of the co-complex: screen. All initial and optimization screening were performed at both 4 °C and 20 °C. Diffracting crystals were obtained from buffer conditions described in Table 1. Crystals for data collection were harvested in the above condition, supplemented with 20 % glycerol (Hampton research, HR2-623) or ethylene glycol (Hampton research, HR2-621) and flash cooled in liquid nitrogen.

**Table 1:** Crystals of NKp46 obtained throughout the study.

| Screen | Condition |
| --- | --- |
| Structure Screen 1+2 HT | E6 - 1.0 M Lithium sulfate, 0.1 M Tris pH 8.5, 0.01 M nickel (II) chloride hexahydrate |
| Structure Screen 1+2 HT | G4 - 0.1 M Sodium HEPES pH 7.5, 20 % w/v PEG 10,000 |
| SG1 Screen HT | B6 - 0.1 M Sodium HEPES pH 7.5, 20 % w/v PEG 10,000 |
| SG1 Screen HT | C7 - 0.1 M Sodium HEPES pH 7.5, 25 % w/v PEG 3,350 |
| SG1 Screen HT | E9 - 0.1 M Potassium thiocyanate, 30 % w/v PEG 2,000 MME |
| PACT Premier HT | E2 – 0.2 M NaBr, 20% PEG 3,350 |
| PACT Premier HT | E5 – 0.2 M NaNO<sub>3</sub>, 20% PEG 3,350 |

A complete X-ray diffraction dataset was collected from a NKp46 protein / Bicycle peptide 4a co-crystal using synchrotron radiation (beamline i03, Diamond Light Source, UK). A total of 3600 diffraction images were collected exposing the crystal to monochromatic X-ray (λ = 0.9763 Å) for 0.01 s during an oscillation of 0.10° (for each image). Full datasets were processed by the automatic expert system Xia2 (pipeline dials) and the unmerged data file processed in aimless to a resolution of 1.75 Å in space group P61 to acceptable statistics (TABLE II).

The NKp46 / Bicycle peptide 4a co-complex structure was solved by molecular replacement with the program PHASER(33) selecting the chain A in two parts (separate pdb files for each domain) from PDB 1P6F.pdb as the search monomers. A single copy of both monomers was placed with confidence (refined TFZ of 59.7). According to Matthews coefficient 1 molecule of NKp46 was expected in the asymmetric unit (AU, solvent content 44.5 %). The obtained solution from the molecular replacement was visually inspected with COOT(34). The only unaccounted map present was consistent with the shape and dimension of the Bicycle peptide 4a. Dictionaries for the Bicycle peptide 4a were obtained with the program ACEDRG(35) using a 2D-formula of the ligand as input (the latter was drawn using Lidia, a graphical user interface for ligand design). The obtained molecular replacement solution was ad hoc modified to place the Bicycle peptide in COOT and refined with REFMAC5(36,37). Further cycles of model building and refinement were carried out until satisfactory statistics were obtained. The final model was analyzed using MolProbity(38) and deposited to the PDB (accession code to be added here). Data collection and refinement statistics for the deposited structure were tabulated (Table 2) and structural figures were generated using Pymol (http://www.pymol.org).

**Table 2:** Data collection and refinement statistics of X-ray diffraction dataset.

| <u>Data Collection</u> |  |
| --- | --- |
| Synchrotron | Diamond |
|  | i03 |
| Wavelength (Å) | 0.976254 |
| Detector type | Eiger2 XE 16M |
| Temperature (°K) | 100 |
| Oscillation range per frame (°) | 0.1 |
| Overall rotation (°) | 360 |
| Space group | P61 |
| Resolution range (Å) | 64.68-1.75 |
| Unit cell parameters (Å), (°) | 74.688 74.688 75.307 |
|  | 90.000 90.000 120.000 |
| Number of observations | 420657 (11487) |
| Number of unique reflections | 24029 (1311) |
| Multiplicity | 17.5 (8.8) |
| Completeness (spherical %) | 99.6 (99.6) [outer = (99.9)] |
| Mean I/ sigma (I) | 11.5 (0.9) |
| Rmerge (all I+/I-) | 0.125 (1.424) |
| Rmeas (all I+/I-) | 0.129 (1.514) |
| Rpim (all I+/I-) | 0.029 (0.505) |
| <u>Refinement</u> |  |
| R work/ R free | 0.209/0.239 |
| Number of atoms |  |
| Protein | 3223 |
| Ligands/ Ions (EDO, TA9) | 33 |
| Water | 159 |
| RMSD |  |
| Bond lengths (Å) | 0.0080 |
| Bond angles (°) | 1.640 |
| Residues in Ramachandran plot | Amino acids (percentage) |
| Favoured | 181 (95.26) |
| Allowed region | 7 (3.68) |
| Disallowed region | 2 (1.05) |
| Mean B Factors | (Å) |
| Amino Acids -NKp46 chain A | 30.6 |
| Amino Acids- BCY Chain B | 34.5 |
| BCY TA9 Ligand | 61.6 |
| Waters | 41.6 |

\*Values for the highest resolution shell are given in parentheses

Data availability statement: The data generated in this study are available upon request from Bicycle Therapeutics via email at TBD (use medical communications email or IR email). Proteins structure information is available in the protein structure database (pdb) as noted above. Chemical structures and synthetic routes of the active compounds used in this study are available in the Supplemental Materials.

## RESULTS

### Generation and characterization of NKp46-binding bicyclic peptides

We used phage display combined with cyclization around a three-fold symmetric scaffold, based on the original methods described for kallikrein inhibitors(29) to identify bicyclic peptides with high binding affinity and selectivity to soluble human NKp46, referred to in the rest of this report as NKp46 bicyclic peptides. The initial NKp46-binding bicyclic peptides, when synthesized chemically, had a biochemical affinity (K_D_) of ∼250 nM. Through rounds of affinity maturation on phage and further optimization via the incorporation of non-natural amino acids (Figure 1A, Supplemental Section 2A) a lead NKp46 bicyclic peptide was identified with a K_D_ of 3.3 nM against soluble human NKp46 protein. The peptide sequence is shown in Figure 1B and in Supplemental section 2 (compound 4b). This bicyclic peptide did not bind to rodent, dog or cynomolgus monkey NKp46 proteins (Figure S1). It also did not compete with the commercial anti-NKp46 antibody clone 9E2 (Figure S1), which conveniently facilitates the simultaneous detection of NKp46 and binding of bicyclic peptides to NKp46.

**Figure 1.**
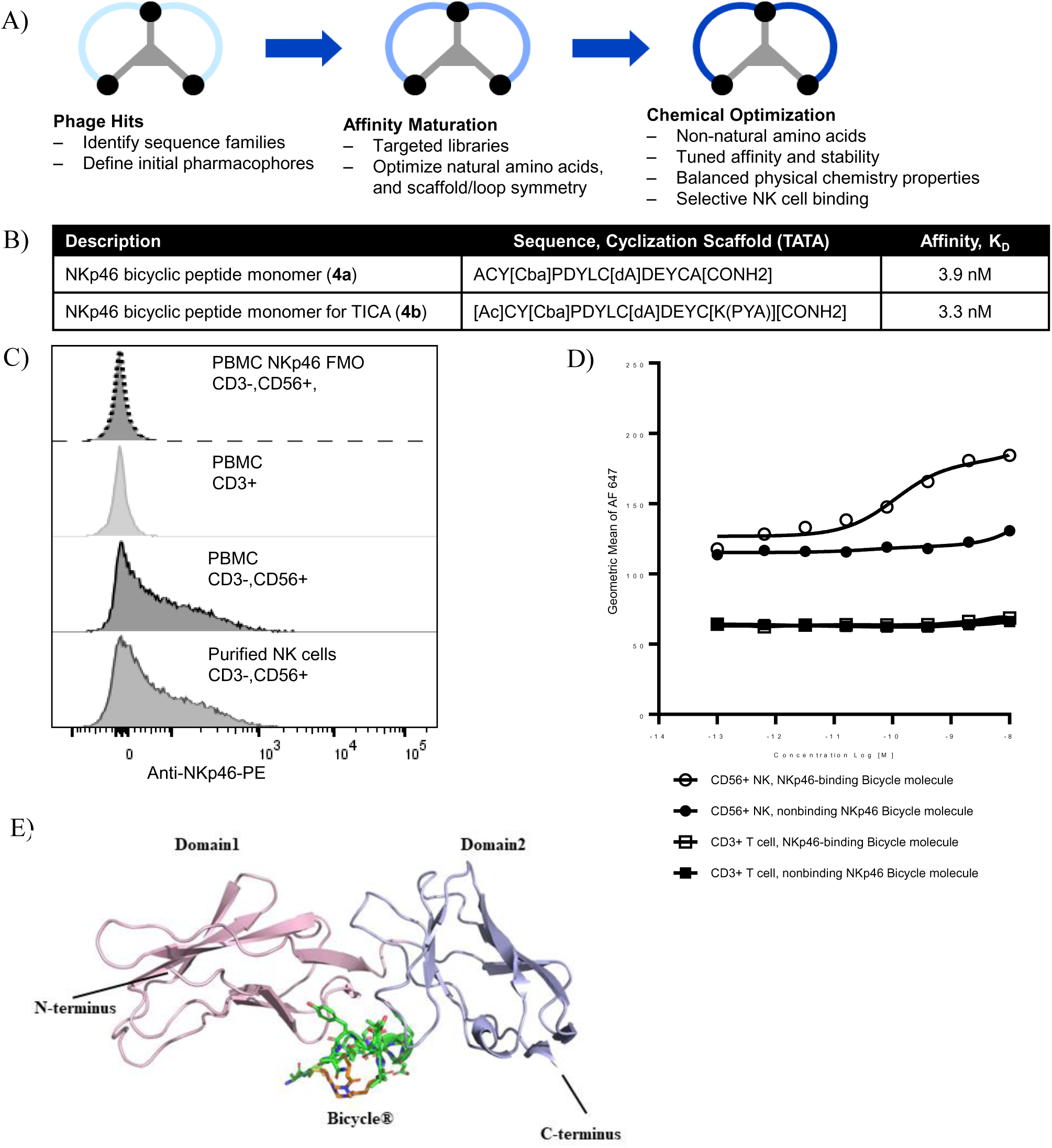
Bicyclic peptides targeting NKp46. A) Schematic representation of the discovery of Bicycle binders using phage display and further modification by chemical synthesis. B) Sequences and biochemical affinities measured using SPR. Linear peptides prepared using Fmoc solid phase chemistry were cyclized using 1,3,5-triacryloylhexahydro-1,3,5-triazine (TATA). Where [Ac] = acetyl, [Cba] = cyclobutyl-L-alanine, [dA] = D-alanine, [K(PYA)] = N-ε-pentynoyl-L-lysine, TATA = 1,3,5-Triacryloylhexahydro-1,3,5-triazine. C) NKp46 Bicycle peptide binds NK cells: NK cells (CD56+, CD16+, CD3-) and T cells (CD3+, CD56-) were characterized for NKp46 expression in PBMC isolated from whole blood (Donor 149) or by negative selection. Shown here is a histogram of a representative PBMC donor with T cells and NK cells. Using anti-NKp46 antibody (Biolegend, clone 9E2), only NK cells were detected to express NKp46. As a control, purified NK cells (by negative selection) were probed with the anti-NKp46 antibody. D) Binding of AF 647-labeled NKp46 Bicycle dimer or NKp46-non-binding Bicycle dimer was monitored in PBMC preparations (shown is NK and T cell populations, other immune cell populations are not shown). Binding of the AF 647-labeled NKp46 Bicycle dimer or NKp46-non-binding Bicycle dimer was monitored in T cells (CD3+ CD56-), NK cells were distinguished by flow cytometry as CD16+ CD56+ CD3-in PBMC preparations. The binding of Bicycle peptides was determined by AF647-signal detection of the cell fraction. Calculated EC_50_ = 39 pM determined by GraphPad Prism®. E) Illustration of NKp46 bicyclic peptide 4a in complex with human NKp46 based on X-ray crystallography data.

We tested binding to cells that endogenously express NKp46, i.e. primary human NK cells. NK cells are characterized by the expression of CD56, CD16, NKp46, and they lack CD3(39)and we used these markers to identify NK cells in preparations of peripheral blood PBMCs (Figure 1C). Fluorescently labeled NKp46 bicyclic peptide dimer (AF647-dimer, compound 13, Supplemental Section 2) bound only to NK cells in the purified PBMCs, with dose-dependency (EC_50_ = 62 pM, average n=3) (Figure 1D). The matching AF647-labeled enantiomeric NKp46 bicyclic peptide dimer did not bind to NK cells. No binding was detected on other lymphocyte populations for either compound (Figure 1D, CD3+ T cell populations and data not shown).

The binding site of the NKp46 bicyclic peptide on human NKp46 protein was elucidated via x-ray crystallography using the bicyclic peptide compound 4a whose peptide sequence is shown in Figure 1B. The binding site sits at the interface between domain 1 (residues 1-100) and domain 2 (residues 100-190) (Figure 1E). The following NKp46 residues from domain 1 are involved in the interaction: Trp-32, Ala-33, Pro-35, His-36, Phe-37, Met-38, Pro-40, Lys-43, Gln-44, Leu-113, Thr-118 and Glu-119. The following residues from domain 2 are involved in the interaction: Leu-156, Glu-158, Gly-159, Arg160, Ser-161, Ser-162, His-163, Arg-189, Phe-191 and Pro-202. The overall structure of NKp46 protein bound to the NKp46 bicyclic peptide remains largely unchanged when compared to apo-NKp46 protein (RMSD of 1.5 Å over 172 Cα, PDB code 1OLL.pdb) (Figure 1E). Several ligands have been identified for NKp46, including ecto-calreticulin, complement factor P (CFP, also known as properdin), vimentin, and viral hemagglutinin(20,21,40,41) (Santara 2023; Narni-Mancinelli, 2017; Garg, 2006; Mandelboim, 2001). In biochemical experiments, the binding of calreticulin was not blocked by NKp46 bicyclic peptides, although the binding of calreticulin was weak (data not shown). We were unable to measure binding of vimentin or CFP to NKp46 protein (data not shown) and so no blocking studies were conducted. Hemagglutinin is reported to bind solely to the membrane proximal domain(42), which is distinct from the binding epitope for the NKp46 bicyclic peptides described in this report, and we did not investigate this further.

### Generation of and functional characterization of conditionally active NKp46 NK-TICA molecules

Building on prior experience with CD137 TICAs, we selected EphA2 as a prototypical tumor antigen that was well characterized and for which we had previously identified EphA2 bicyclic peptides that were competent to scaffold immune receptor-binding bicyclic peptides converting them into conditionally functional agonists(30). Briefly, EphA2 is a receptor tyrosine kinase that is expressed on a range of epithelial tumors and whose expression has been correlated with disease progression and poor prognosis(43) and references therein. The same EphA2 bicyclic peptide as used in our prior work (biochemical K_D_ ∼2 nM) was conjugated to two of the NKp46 bicyclic peptide (4b) via polyethylene glycol (PEG)-based linkers to create bispecific NK-TICA molecules (Figure 2A and Supplemental Section 2B). This Bicycle molecule then has a 1:2 format. We also synthesized corresponding NK-TICA molecules where either the EphA2 bicyclic peptide or the NKp46 bicyclic peptide were replaced with an enantiomeric bicyclic peptide having the same amino acid sequence but with no affinity for the respective target (Supplemental Section 2B). We used these as control molecules to dissect the specificity and conditional nature of NK-TICA molecules.

**Figure 2.**
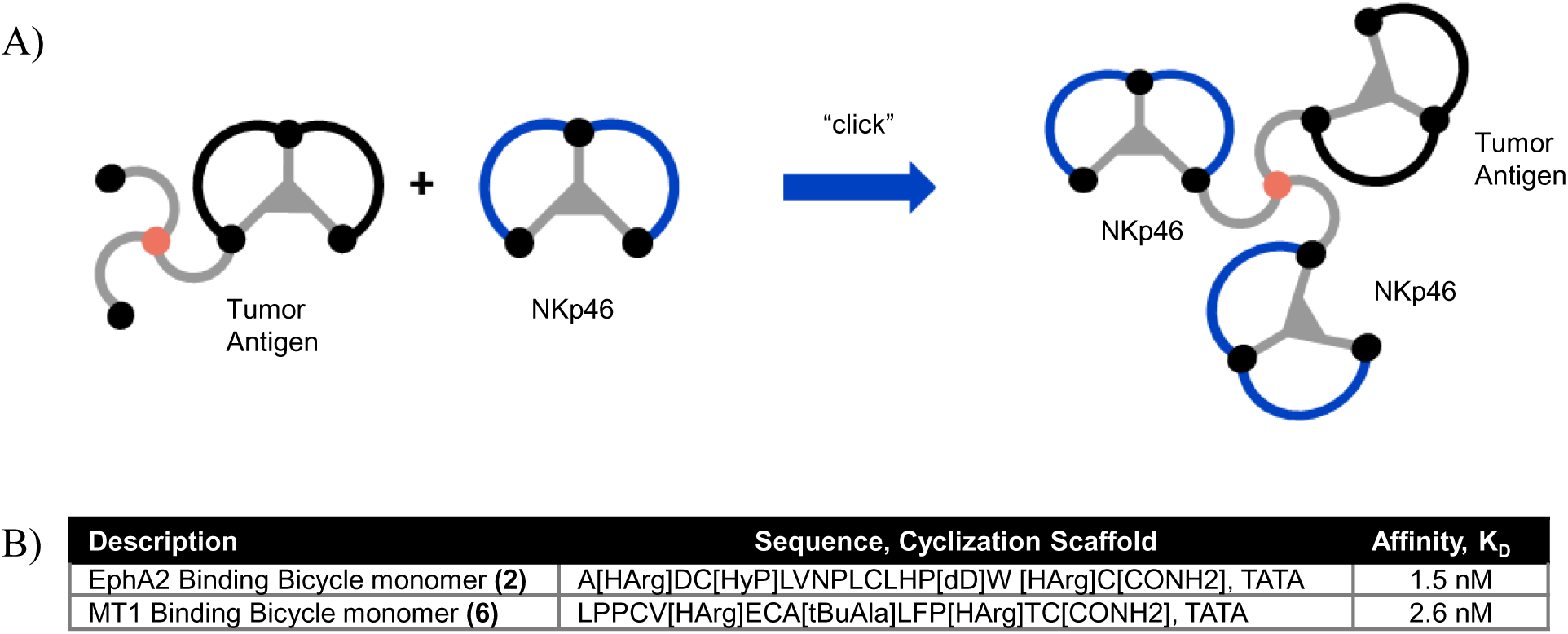
Bispecific NK-TICA molecules. A) Schematic diagram for the synthesis of EphA2/NKp46 and MT1/NKp46 NK-TICA molecules. As previously reported for CD137 and illustrated here, PEG’ylated tumor antigen bicyclic peptides were combined with NKp46 bicyclic peptide through copper catalyzed “click” chemistry to generate a 1:2 format NK TICA molecules. B) Sequences and biochemical affinities measured using SPR. Linear peptides prepared using Fmoc solid phase chemistry were cyclized using 1,3,5-triacryloylhexahydro-1,3,5-triazine (TATA), where [Ac] = acetyl, [dA] = D-alanine, [dD] = D-aspartic acid, [HyP] = trans-4-hydroxy-L-proline, [Harg] = L-homoarginine, [K(PYA)] = N-ε-pentynoyl-L-lysine, [tBuAla] = t-butyl-L-alanine, [Cha] = cyclohexyl-L-alanine, [Agb] = L-norarginine, [2FTyr] = 2-fluoro-L-tyrosine, [6FTrp] = 6-fluoro-L-tryptophan, [NMeAla] = N-methyl-L-alanine, [dK] = D-lysine, TATA = 1,3,5-tripropanoylhexahydro-1,3,5-triazine.

As described in the Methods section, HT1080 or A431 cells that express recombinant luciferase were cultured and exposed to primary human NK cells. A reduction in measured luminescence equates proportionally to fewer live tumor cells in the culture. Addition of EphA2/NKp46 NK-TICA molecules to the culture caused a dose-dependent increase in killing of HT1080 cells by NK cells (Figure 3A, EC_50_ = 0.5 pM). Control NK-TICA molecules (non-binding for NKp46 or non-binding for EphA2) were inactive in this assay (Figure 3A) therefore. As with CD137 TICAs, the functional activation of NK cells by NK-TICAs was conditional upon binding of the tumor antigen as well as NKp46. We also prepared 1:1 format NK-TICA molecules, which were similarly active but less potent (data not shown). We did not make or test the corresponding 1:3 format. The EphA2/NKp46 NK-TICA molecules also directed NK killing of A431-luc cells (Figure 3C). There was no evidence of fratricide because NK cell viability was similar in the presence of binding and non-binding NK-TICA molecules at 24 hr (Figure 3B). We also prepared and tested NK-TICA molecules using bicyclic peptides that bind to the tumor-associated membrane type 1 matrix metalloproteinase MT1-MMP (MT1). Briefly, MT1 is of interest as a targeting antigen due to overexpression in important epithelial cancers and association with prognosis, and we have described previously bicyclic peptides that bind to MT-1(32) and references therein. The NKp46 bicyclic peptide 4b was conjugated to an MT1 bicyclic peptide (K_D_ ∼15 nM) (Figure 3C and Supplemental Section 2) to again create a 1:2 format NK-TICA molecule. Like EphA2, MT-1 proved competent to scaffold NKp46 bicyclic peptides in a manner that afforded potent and conditional NK cell activation in co-culture with MT1-expressing tumor cells, in this case HT1080-luc cells (Figure 3D). Thus, the function of NK-TICA molecules is not limited to a single antigen or a single tumor cell line.

**Figure 3.**
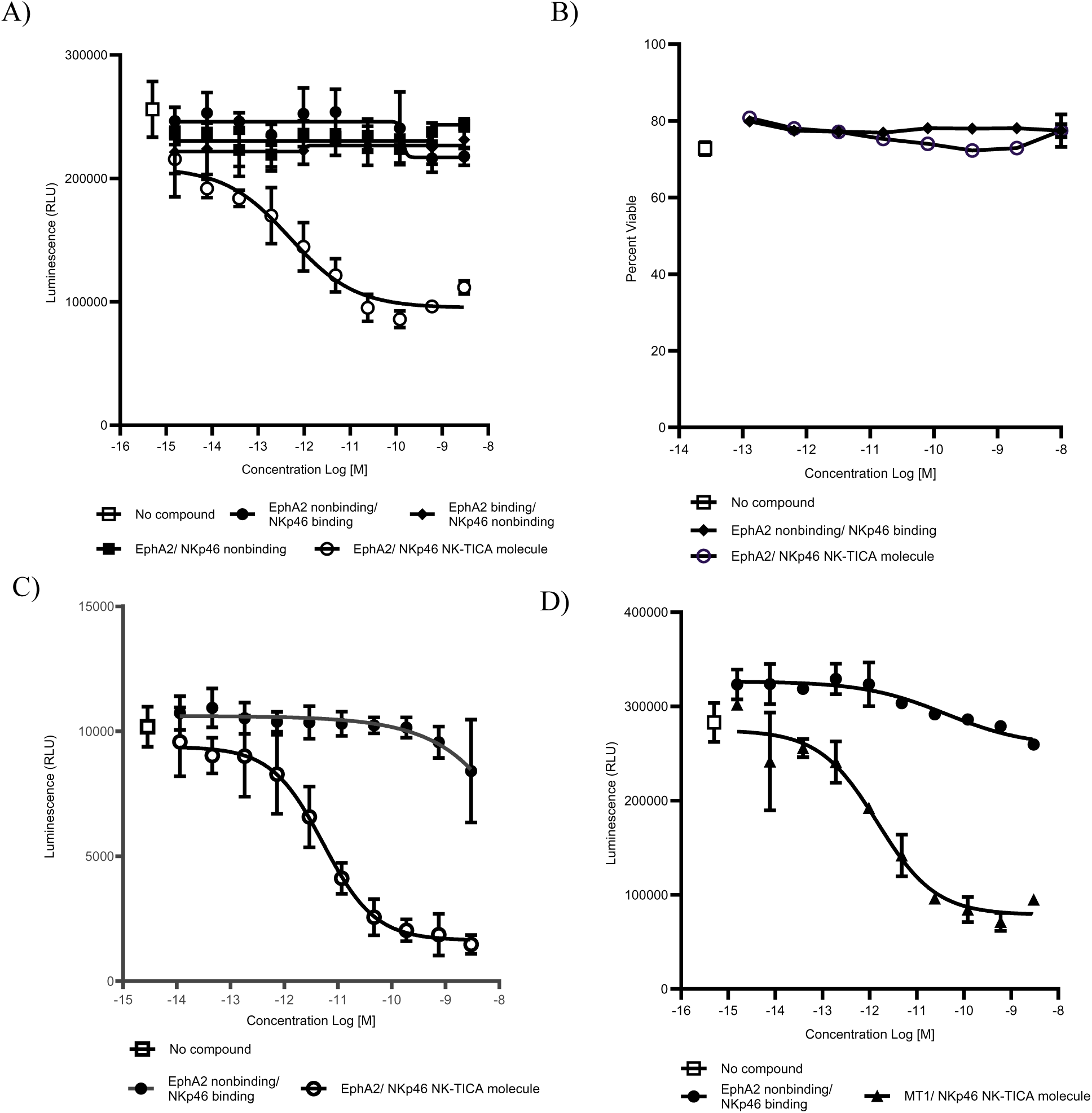
NK TICA molecules drive tumor cell killing by primary human NK cells. A) NK cells specifically kill HT1080-luc cell (E:T ratio 5:1) in the presence of NK-TICA® molecule bearing NKp46-binding and EphA2-binding Bicycle monomers (EC_50_ = 0.5 pM). Without EphA2 binding, alone or in combination with non-binding NKp46, no enhanced NK-mediated killing of HT1080 cells was observed (non-binding NK-TICA molecules). B) Cells in co-culture were stained with viability dye and anti-CD56 antibody for detection of viable NK cells. The percent NK cells were gated for incorporation of viability stain and plotted as percent viable CD56+. C) NK killing of A431-luc cells (E:T ratio 10:1) was demonstrated with the EphA2-binding NK-TICA molecule, EC_50_ = 5 pM. D) NK killing of HT1080-luc cells in the presence of MT1/NKp46 NK-TICA molecules, EC_50_ = 8 pM. In each panel, luminescence values for tests in the absence of compound were arbitrarily shown at 5 x 10^-15^ M.

### The NK-TICA molecule elicits conditional NK cell activation and cytokine production in the presence of tumor cells

Upon recognition of foreign or transformed cells, NK cells can release proteases, cytokines, and chemokines, such as granzyme B, interferon-gamma (IFN-γ), and fms-3-like ligand (FLT3LG) in a controlled and directed manner(1). Thus, we explored the ability of the EphA2/NKp46 NK-TICA molecule to enhance these aspects of NK cell function. NK cells in co-culture with HT1080-luc cells secreted TNFα (EC_50_ =15 pM) and IFNγ (EC_50_ =15 pM) at 4 hr and FLT3LG (EC_50_ = 6 pM) at 48 hr in the presence of NK-TICA® molecules (Figure 4A TNFα, 4B INFγ, 4C FLT3LG). Additionally, the NK-TICA molecules required competent NKp46-binding and EphA2-binding bicyclic peptide arms to induce cytokine production, as the non-binding NK-TICA analogs did not induce measurable cytokine secretion. CD25, the high affinity IL-2 receptor alpha subunit, can be expressed on activated NK cells, with a putative role in NK cell survival to promote capture of interleukin-2 for induction of transcription and translation of cytotoxic granules(4). Increases in CD25 expression were detected in a dose-dependent manner on NK cells treated with NK-TICA molecules, at 24 hr in co-culture with HT1080-luc cells (EC_50_ = 7 pM) (Figure 4D).

**Figure 4.**
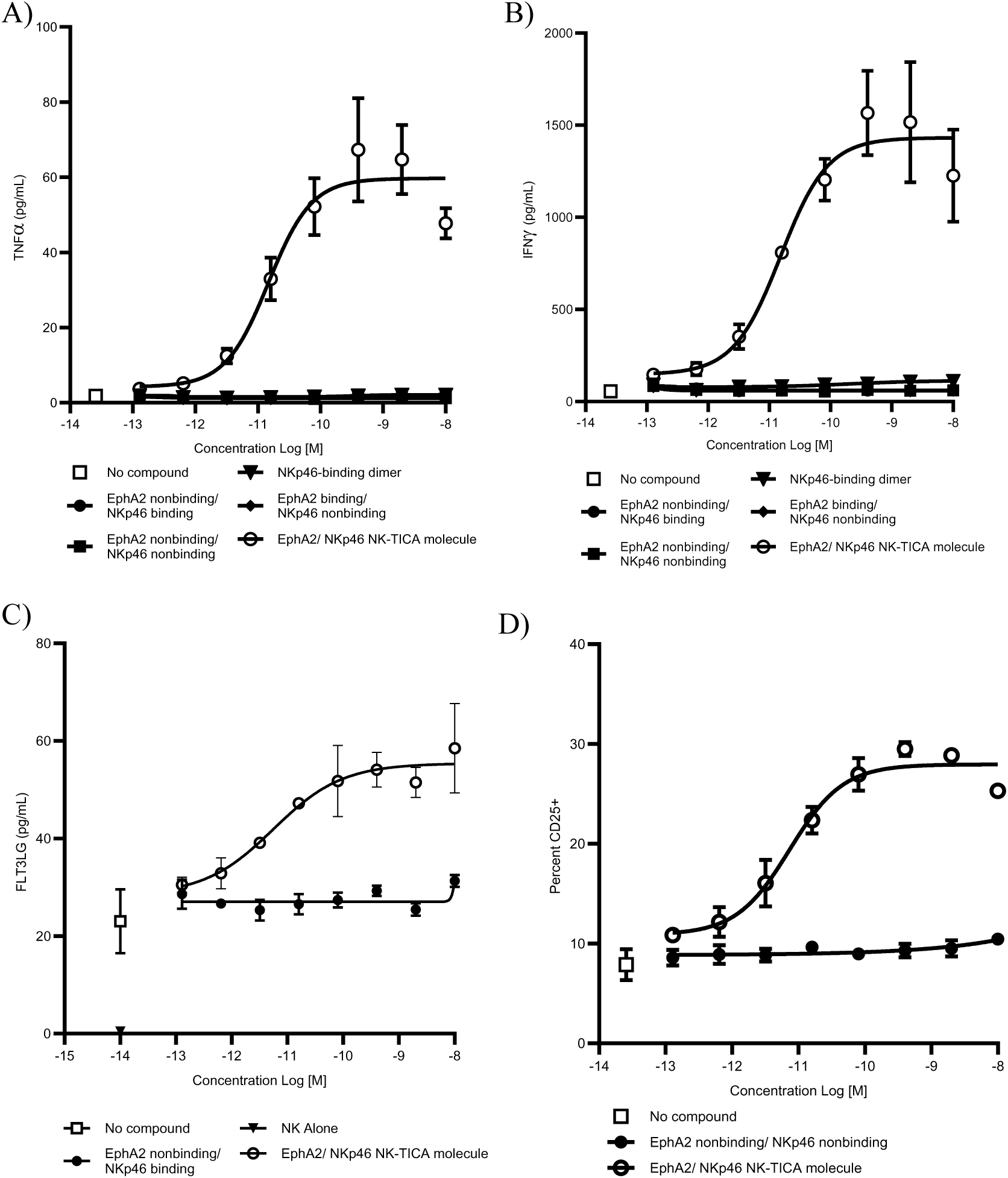
Induced cytokine secretion and CD25 expression by NK cells treated with NK-TICA molecules. NK cells were co-cultured with HT1080-luc (E:T ratio 5:1) with the addition of the NK-TICA molecules. A&B) TNFα (A) and IFNγ (B) released into supernatants after 4 hr were measured by MSD® cytokine kit. C) Supernatant FLT3 ligand after 48 hr of co-culture. D) Cells in co-culture were stained with viability dye, anti-CD56 antibody and anti-CD25 antibody for detection of viable CD56+ CD25+ NK cells, measured by flow cytometry. The percent viable CD56+ CD25+ NK cells were plotted as percent viable CD56+, CD25+ NK cells after 24 hr in co-culture conditions.

Interestingly, this increase was limited to ∼20% of the total population of NK cells in the culture, suggesting some heterogeneity within the NK cells purified from PBMCs for these experiments. No increase in CD25 expression on NK cells was detected in conditions including the non-binding NK-TICA molecules.

### The NK-TICA molecule promotes NKp46 clustering and localization of lysosomal granules to the NK:tumor cell interface

Previous reports have suggested the need of NKp46 clustering to the immune cell-tumor cell interface for NKp46-dependent NK killing of tumor cells with immune synapse formation characterized by NKp46 receptor clustering as well as the sequestration of lytic vesicles for the release of cytotoxic contents(44,45). We explored the ability of EphA2/NKp46 NK-TICA molecules to enhance the NK cell interaction with HT1080 cells through time-lapse confocal microscopy. Initially, we detected the expression of NKp46 on NK cells from two donors with an anti-NKp46 antibody (red, Alexa Fluor® 647) (Figure 5A). As shown in Figure 5, NKp46 (red) was displayed on NK cells with a uniform distribution on the surface when NK cells were plated in co-culture with HT1080 cells (green) (Figure 5A No NK-TICA molecule). With the addition of NK-TICA molecule (Figure 5B), the detected NKp46 distribution shifted to a more localized pattern, whereby NKp46 and intracellular granules (labeled with LysoView™) (blue) became punctate and localized to the NK cell – tumor cell interface (Figure 5B: NKp46 stained with anti-NKp46-Alexa Fluor® 647 antibody (red)). This suggests that the NK-TICA molecules can cluster the NKp46 receptor and sequester cytotoxic granules for the generation of an immune synapse, resembling data reported previously(44,45). Next, we constructed an AF647-conjugated NK-TICA molecule to allow us to monitor the NK-TICA molecule and its distribution on the NK cell surface in a co-culture condition (Supplemental section 2). This AF647-conjugated NK-TICA molecule displayed similar distribution on the cell surface (Figure 5C) as for the NKp46 antibody. Addition of either the AF647-conjugated NK-TICA molecule (Figure 5C) or unlabeled NK-TICA molecule combined with anti-NKp46 Alexa Fluor® 647 antibody (Figure 5B) displayed clustering of the NKp46 receptor and localization of lytic vesicles to the tumor interface. Addition of the NK-TICA molecule with the non-binding EphA2 Bicycle monomer resulted in surface distribution of NKp46 that was similar to addition of no NK-TICA molecule added (Figure 5A, D). Therefore, we confirm the requirement of both tumor antigen and NKp46 binding for the demonstration of NK-tumor synapse formation.

**Figure 5.**
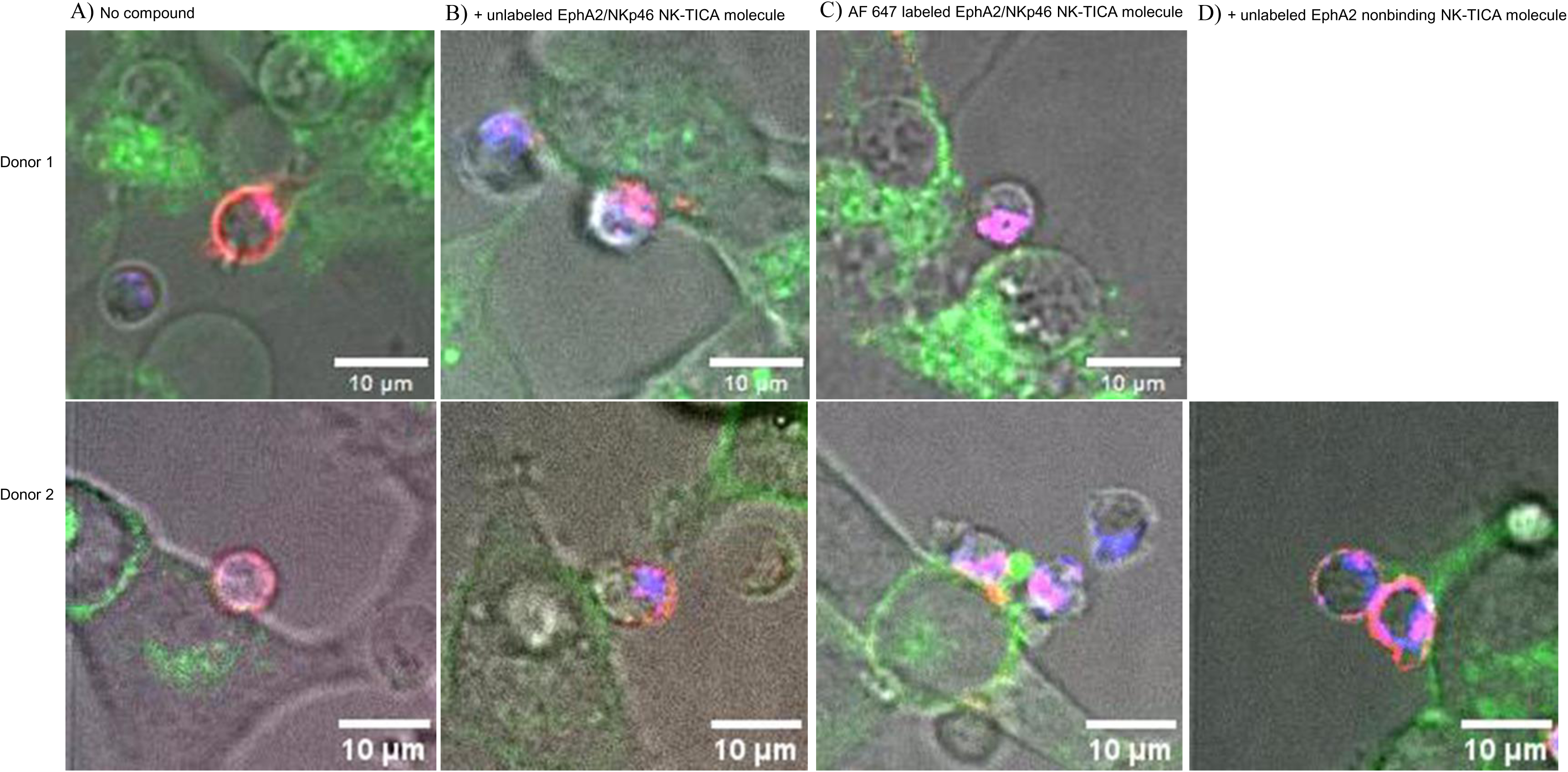
NK-TICA molecules enhance synapse formation at the NK-tumor cell interface. Two donors NK cells were labeled with LysoView™ 594 (Blue channel) in co-culture with HT1080 cells (Green channel; CellBrite®™ Steady 488 membrane dye Biotium®) (Top: Donor 1, Bottom: Donor 2). In A, B, D, NKp46 was stained with anti-NKp46 Alexa Fluor® 647 antibody (Red channel). A) NK in co-culture with HT1080 cells. B) NK in co-culture with HT1080 with addition of NK-TICA molecule (2 nM). C) NK in co-culture with HT1080 cells with addition of AF 647 labeled NK-TICA molecule (2 nM). D) NK in co-culture with HT1080 cells with addition of EphA2 non-binding NK-TICA molecule (2 nM).

### NK cells are capable of successive rounds of enhanced killing of tumor cells in the presence of the NK-TICA molecule

We next evaluated the ability of NK cells treated with the NK-TICA molecule to target successive rounds of tumor cells. For the maintenance of viability and regeneration of cytolytic granules, NK cells were pretreated with interleukin 2 (IL-2). Labeled HT1080-Nuc Light Green cells were plated on a flat-bottom tissue culture plate to permit detection on an Incucyte® S3 reader. NK cells were added to the culture wells containing the adherent tumor cells at an E:T ratio of 10:1 in the presence or absence of NK-TICA molecules (Figure 6). In order to visualize and measure a reduction in HT1080-Nuc Light cells, the NK cells were permitted to settle to the bottom of the culture plate wells and a loss of green fluorescence detection was measured over a time span of 24 hr. After 24 hr of co-culture, the NK cells eliminated the HT1080 cells from the wells, regardless of the presence of the NK-TICA molecule. A second round of HT1080-Nuc Light tumor cells was added to each culture well once we removed half of the volume from the top of each well. This permitted us to challenge the NK cells without adding more NK-TICA molecules or NK cells. This was repeated over the time span of 72 hr for three rounds of live HT1080-Nuc Light cells addition. In each round there was a drop in the number of tumor cells (Figure 6). Upon second exposure to tumor cells, the NK cells that had received prior the NKp46-binding and EphA2-binding NK-TICA molecule resulted in 88% tumor killing, compared to 63% with the control compound (NKp46-binding EphA2-nonbinding NK-TICA molecule). It is noteworthy that the IL-2-pretreated NK cells with the NK-TICA molecule demonstrated more efficient killing of the tumor in the third round of co-culture, at 72 hr (NK-TICA molecule). Thus EphA2/NKp46 NK-TICA molecules could enable sustained killing of tumor cells over time and over repeated exposure to fresh tumor cells.

**Figure 6.**
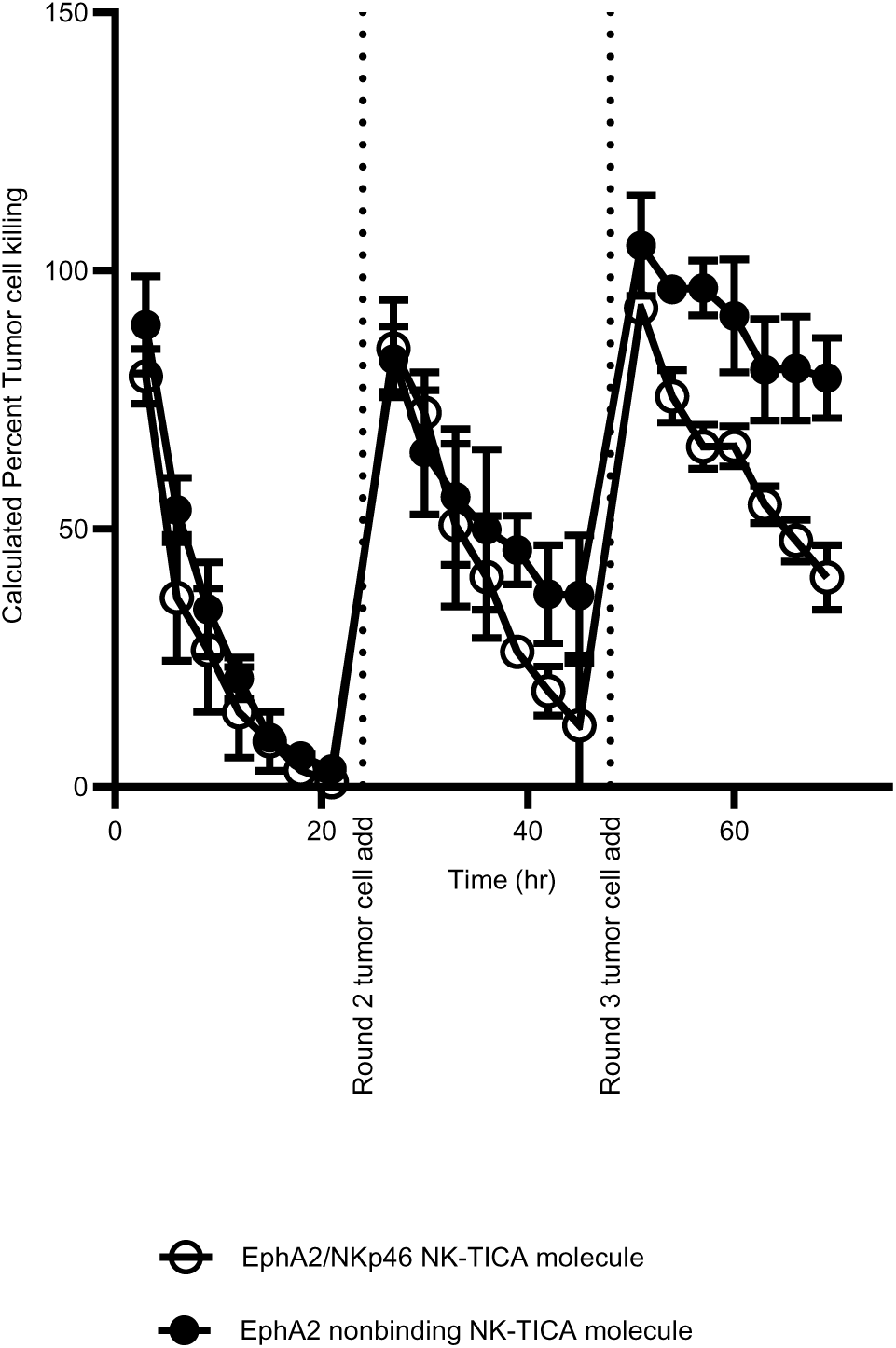
NK-TICA molecules enable sustained serial tumor cell killing by primary human NK cells. IL-2-stimulated NK cells (20000) in co-culture with HT1080 IncuCyte® NucLight Green target cells (GFP-positive, 2000) (10:1 E:T ratio) in the presence of 2 nM NK-TICA molecule (open circle) or control NKp46-binding EphA2-nonbinding NK-TICA molecule (closed circle) (n=3). Tumor cells (2000) were added at a 10:1 effector-to-target cell ratio every 24 hr for 3 rounds. GFP monitoring was captured by IncuCyte® over 120 hr. The calculated percent loss of GFP was determined by seeding GFP intensity equaling 100% and subtraction of each well GFP intensity at each time point.

## DISCUSSION

In this report we describe the discovery of chemically synthetic, small, constrained bicyclic peptides that can bind to human NKp46 both biochemically and on the surface of primary human NK cells, and we report on the structural basis for that interaction (Fig. 1). We were then able to conjugate the NKp46 binders to other bicyclic peptides that bind to tumor antigens, exemplified in this work by EphA2 and MT-1, in such a way as to afford conditional activation of the killing function of NK cells (Figs. 2 & 3). We called this class of molecules NK tumor immune cells agonists (NK-TICAs). As we found with CD137 TICAs in the same 1:2 format(31), the activation of immune receptors was completely conditional on binding to tumor antigen. This absolute dependence on tumor antigen, combined with NK-TICAs being small synthetic chemicals with no propensity for aggregation, denaturation or oligomerization, mean that they are predicted to be inert with respect to immune cell activation in the absence of antigen (i.e. in the peripheral blood). We believe this is a critically important feature that discriminates Bicycle TICAs from antibody- or protein-based immune agonists. It is well known that, in vitro, and indeed in vivo in preclinical mouse models, antibodies can activate specific NK cell killing of tumor cells. We found that the extent of killing in the presence of NK-TICAs was similar to that triggered by anti-EphA2 antibodies, presumably acting through CD16 (data not shown). We did not have a suitable anti-MT-1 antibody available, but the same qualitative finding of similar extent of kill was true for a third proprietary tumor antigen, not disclosed in this report, in comparison to the corresponding clinical antibody (data not shown).

NK cells treated with NK-TICAs in the presence of tumors cells also secreted pro-inflammatory cytokines consistent with the concept that NK cells sit at the interface of innate and adaptive tumor immunity (Fig. 4). This included production of FLT3L which is an essential factor for recruitment and activation of conventional dendritic cells that are central to development of a productive anti-tumor immune response(5,46,47). We also provide qualitative confocal imaging data to visualize the interaction of NK cells with tumor cells in the presence of NK-TICAs and the effect of those compounds on the distribution of NKp46 on the cell surface (Fig. 5). Remarkably, NK-TICAs could also prolong the killing function of NK cells through multiple re-additions of tumor cells to the culture suggesting that NK-TICAs may mitigate the exhaustion of NK cells over repeated round of killing (Fig. 6). This last observation is highly germane given the assertion in the field that intratumoral NK cells are dysfunctional or exhausted(11,48).

Important questions that are not addressed fully in this report include whether the NK-TICAs or more the NKp46-binding component can block or otherwise modulate the activation of NKp46 by host or pathogen ligands. As noted in the results section, our investigations in this regard yielded equivocal results and more work is needed to fully understand both any safety liabilities from interrupting the normal function of NKp46, but also the potential for NKp46 blocking agents as therapeutics in inflammatory disease settings. Another unresolved question is whether the activation of NK cells (killing, cytokine secretion) arises from proximity to tumor cells, stabilization of an immune synapse facilitating sensing of stress signals, activation of NKp46 signaling per se, or some combination of each.

We have not investigated the molecular signaling downstream of NKp46 in our models, but this remains a question of high interest. Finally, we have not addressed whether NK-TICAs can promote tumor rejection in vivo. We evaluated two humanized mouse models because the NKp46 binders used in this report do not bind to mouse NKp46. First, immune system-humanized mice bearing A431 and HT1080 subcutaneous xenografts or A549 experimental lung metastases, and second, mice transgenic for human NKp46. However, we found that both types of model lacking in important aspects, for example, lack of NK cell infiltration into the tumors in humanized mice(49).

In summary, we have created the first chemically synthetic, small molecule, conditionally active NK cell engagers, termed NK-TICAs, and demonstrated their activity in the setting of primary human NK cells killing human tumor cells in vitro. This work represents an important step towards testing the concept that NK cells can be harnessed to drive eradication of tumors in cancer patients. The work builds on success with CD137-engaging TICAs, opening a door to a new general approach for therapeutic activation of immune cell receptors.

## SUPPLEMENTAL MATERIALS

### Supplemental Section 1 – Biochemistry SPR - Methods

#### Surface plasmon resonance

For analysis of monomer binding, Biacore™ 8k+ instrument was used with a CM5 chip (Cytiva™). SPR experiments were performed to determine k_a_ (M-1s-1), k_d_ (s-1), K_D_ (nM) values, where measurable, of Bicycle monomers against immobilized NKp46 (Fc-tagged human NKp46 (ACROBiosystems catalog # NC1-H5257 and C365P1-202KF1-S3), EphA2 (1), and MT1 (1).

All experiments were carried out at 25 °C. Anti-human IgG (Fc) antibody (Cytiva) was immobilized on all flow cells by standard amine coupling chemistry. The running buffer was HBS-N (Cytiva, 10 mM HEPES, 0.15 M NaCl, pH 7.4) and the flow rate was 10 µL/min. The carboxymethyl dextran surface was activated with a 1:1 ratio of 0.4 M 1-ethyl-3-(3-dimethylaminopropyl) carbodiimide hydrochloride (EDC)/0.1 M N-hydroxy succinimide (NHS) (Cytiva) for 420 s. Anti-human IgG (Fc) antibody was diluted to 12.5 ug/mL in 10 mM sodium acetate pH 5.0 and injected onto the chip surface for 360 s. Residual activated groups were blocked with a 420 s injection of 1 M ethanolamine (pH 8.5). The final surface density was 3000 – 5000 RU. Fc-tagged human NKp46 was diluted to 0.1 uM in HBS-N and was captured on flow cell 2 only of the sensor chip at a flow rate of 10 µL/min to 500 – 1000 RU. The buffer was then changed to PBS-P+ (Cytiva) supplemented with 2 % DMSO. Peptide dilutions were prepared in this buffer to give a final concentration of 2 % DMSO. The flow rate was 50 µL/min with 60 s association and 400 s dissociation. Data was corrected for DMSO volume exclusion effects. All data were double referenced against blank injections and reference surface response using standard processing procedures. Data processing and kinetic fitting was performed using Biacore™ Insight Evaluation Software with Extended Screening and Characterization Extension (Cytiva). Data were fitted using steady state affinity 1:1 binding model to determine K_D_. Data were fitted using kinetic 1:1 binding model to determine kinetic constants where appropriate. The NKp46 bicyclic peptide 4b was also tested using the SPR techniques described above on other species NKp46 proteins: rodent, rat/mouse (GenScript catalog # U851GHC310_3), canine (GenScript catalog # U851GHC310_3), monkey, cynomolgus macaque (ACROBiosystems catalog # NC1-C52H4).

**Supplemental Table 1:** Sources of commercial antibodies used in this work.

| Antibody or reagent | Source, catalog number |
| --- | --- |
| Fixable Viability Dye | eBioscience/ThermoFisher catalog #65-0865-18, 65-0865-14 |
| anti-CD3 | BD Bioscience catalog #612750 |
| anti CD56 | BD Bioscience catalog #740299 |
| anti-CD20 | Biolegend catalog #302332 |
| anti-CD11C | Biolegend catalog #301628 |
| Anti-NKp46 | Biolegend catalog #331908, BD Biosciences catalog #557911 |
| anti CD8 | Biolegend catalog #301046 |
| anti CD14 | Biolegend catalog #982502 |
| Anti-MT1 | R&D systems catalog #FAB9181P |
| Anti- EphA2 | R&D systems catalog #FAB3035P |
| IgG1 isotype | Biolegend catalog #400112 |
| IgG2b isotype | R&D systems catalog #IC0041P |
| Anti-CD16 | Biolegend catalog #302038 |
| Anti-CD25 | Biolegend catalog #302634 |

#### SPR - Results

Affinity was determined by surface plasmon resonance (SPR) against human, rat, dog and monkey (cynomolgus) NKp46 (Fig S1A, C, D, E). No separate experiments for mouse NKp46 were performed because mouse and rat NKp46 are identical in the extracellular domain and full length mouse protein was not available with sufficient quality. For competition experiments the commercial anti-NKp46 antibody (clone 9E2) was purchased from BD Biosciences, catalog #557911. Competition of binding for human NKp46 by anti-NKp46 antibody (clone 9E2) and NKp46 bicyclic peptide 4b was determined by SPR, up to 500 nM, no competition being observed (Fig. S1B). The affinity of MT-1 Bicycle monomer to human MT1 protein was determined by SPR (Fig. S1F).

### Supplemental Section 2 – Chemistry Synthesis of Bicycle monomers

Briefly, linear peptide was synthesized on Rink amide resin using standard Fmoc (9-fluorenylmethyloxycarbonyl) solid phase peptide synthesis, either by manual coupling, or with an automated peptide synthesizer (e.g. Biotage® Syro II), as previously described. The peptides were cleaved from the resin using a TFA cleavage cocktail containing appropriate protecting group scavengers and the peptide were precipitated with diethyl ether and dissolved in 50:50 acetonitrile/water. The crude peptide was cyclized with 1,3,5-Triacryloylhexahydro-1,3,5-triazine (TATA) or 1,3,5-(tri-2-bromoacetyl)-hexahydro-1,3,5-triazine at ∼1 mM concentration peptide with 1.3 equivalents scaffold, using ammonium bicarbonate (100 mM) as base. Cyclization was monitored by matrix-assisted laser desorption ionization time-of-flight (MALDI-TOF) or LC−MS. Once cyclization was complete, the reaction was quenched with N-acetyl cysteine (10 equivalents). The solutions were lyophilized and purified by RP-HPLC. Fractions containing the desired product at sufficient purity and correct molecular weight (verified by either MALDI-TOF and HPLC, or LC-MS) were pooled and lyophilized.

### Synthesis of NK-TICA molecules

#### For the synthesis of Bicycle intermediates and TICAs

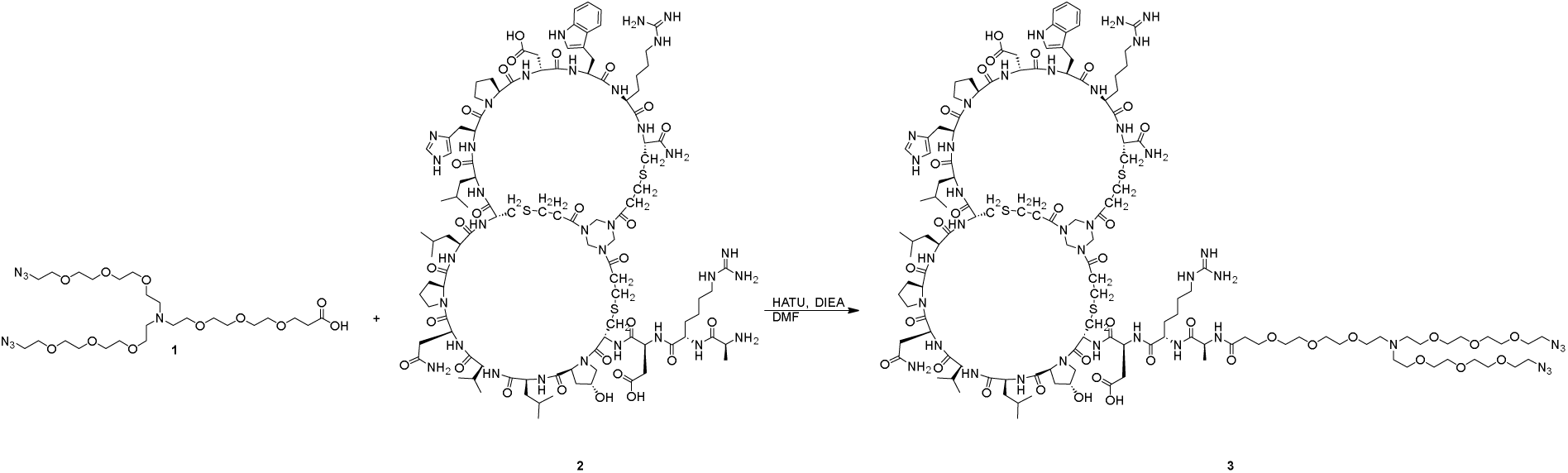

Bicycle peptide intermediate, 3, was synthesized as previously described (Upadhyaya P et al. 2021). To a mixture of compound 1 (60.0 mg, 96.2 μmol, 1.0 eq.) in DMF (3 mL) was added DIEA (12.4 mg, 96.2 μmol, 16.8 μL, 1.0 eq.) and HATU (38.4 mg, 101 μmol, 1.05 eq.) and the mixture stirred for 5 min. Then EphA2-binding Bicycle, 2, (243 mg, 101 μmol, 1.05 eq.) was added to the mixture, which was purged with N_2_, then stirred at 40 °C for 16 hr under N_2_ atmosphere. LC-MS showed compound 1 was consumed completely and one main peak with desired m/z was detected. The reaction mixture was purified by preparative-HPLC to give 3 (154 mg, 48.1 μmol, 50.0 % yield, 94.0% purity) as a white solid. Calculated MW: 3006.48, observed m/z: 1002.8 ([M+3H]^3+^), 1504.4 ([M+2H]^2+^).

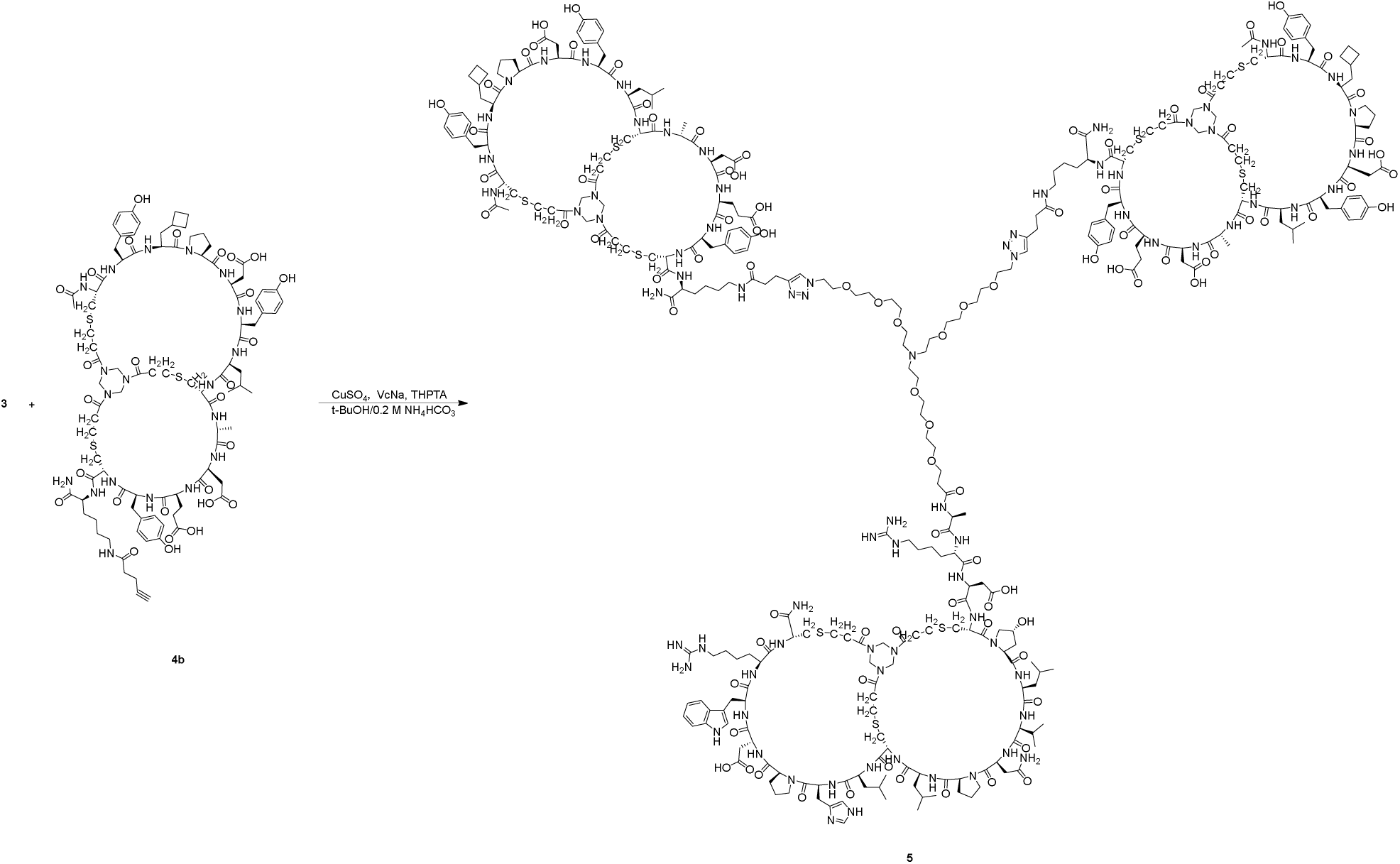

A mixture of 3 (20.0 mg, 6.65 µmol, 1.0 eq.), NKp46-binding Bicycle monomer 4b, (30.5 mg, 14.64 µmol, 2.2 eq.), and THPTA (5.8 mg, 13.30 µmol, 2.0 eq.) was dissolved in t-BuOH/H_2_O (1:1, 1.0 mL, pre-degassed and purged with N_2_ for 3 times), and then aqueous solution of CuSO_4_ (0.4 M, 24.9 µL, 1.5 eq.) and sodium ascorbate (VcNa) (4.0 mg, 19.96 µmol, 3.0 eq.) were added under N_2_ atmosphere. The pH of this solution was adjusted to 8 by dropwise addition of 0.2 M NH_4_HCO_3_ (in 1:1 t-BuOH/H_2_O), and the solution turned to light yellow. The reaction mixture was stirred at 25-30 °C for 0.5 hr under N_2_ atmosphere. LC-MS showed compound 4b consumed completely, and one main peak containing desired m/z (calculated MW: 7169.18, observed m/z: 1434.9 ([M+5H]^5+^), 1195.9 ([M+6H]^6+^)) was detected. The reaction mixture was filtered to remove the undissolved residue. The crude product was purified by prep-HPLC, and 5 (24.0 mg, 3.35 µmol, 50.32 % yield. 97.0 % purity) was obtained as a white solid.

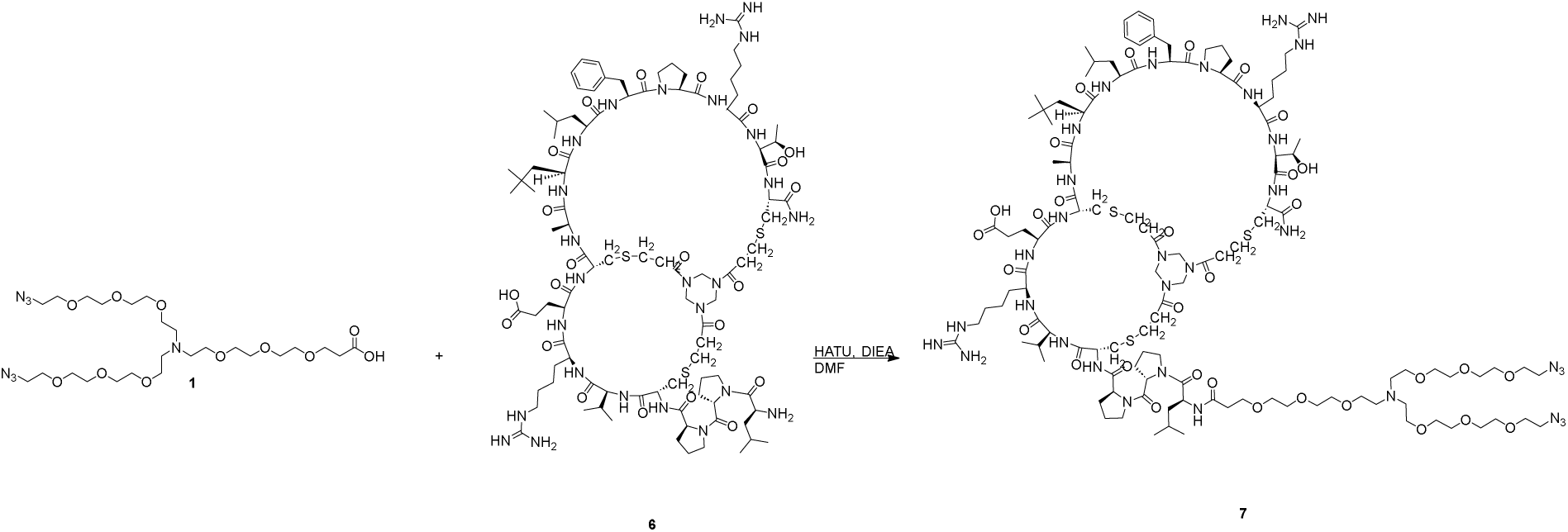

A mixture of compound 1 (20.0 mg, 32.1 µmol, 1.0 eq.), HATU (13.4 mg, 35.3 µmol, 1.1 eq.) and DIEA (12.4 mg, 96.2 µmol, 16.8 µL, 3.0 eq.) was dissolved in DMF (1.0 mL). The reaction mixture was activated at 25-30 °C for 6 min, then MT1-binding Bicycle monomer 6, (74.4 mg, 35.3 µmol, 1.1 eq.) was added into the reaction mixture. The reaction mixture was stirred at 25-30 °C for 0.5 hr. LC-MS showed one main peak with desired m/z (MW: 2714.27, observed m/z: 1358.1 [M+2H]^2+^, 905.9 [M+3H]^3+^, 679.5 [M+4H]^4+^) was detected. The reaction mixture was filtered to remove the undissolved residue. The residue was then purified by prep-HPLC. Fractions containing the desired product were pooled, frozen, and lyophilized to give 7 (42.3 mg, 15.6 µmol, 48.5 % yield, 97.7 % purity) as a white solid.

A mixture of 7 (15.0 mg, 5.53 µmol, 1.0 eq.), and Bicycle monomer 4b, (24.2 mg, 11.6 umol, 2.1 eq.) and THPTA (2.4 mg, 5.53 µmol, 1.0 eq.) was dissolved in t-BuOH/H_2_O (1:1, 2.0 mL, pre-degassed and purged with N_2_ for 3 times), and then aqueous solution of CuSO_4_ (0.4 M, 20.7 uL, 1.5 eq.) and sodium ascorbate (VcNa) (3.3 mg, 16.6 µmol, 3.0 eq.) were added under N_2_. The pH of this solution was adjusted to 8 by dropwise addition of 0.2 M NH_4_HCO_3_ (in 1:1 t-BuOH/H_2_O), and the solution turned to light yellow. The reaction mixture was stirred at 25-30°C for 1 hr under N_2_ atmosphere. LC-MS showed 4b was consumed completely, and one main peak with desired m/z (calculated MW: 6876.9, observed m/z: 1376.4 [M+5H]^5+^, 1147.3 [M+6H]^6+^) was detected. The reaction mixture was filtered to remove the insoluble residue. The crude product was purified by prep-HPLC, and 9 (19.1 mg, 2.77 µmol, 50.0 % yield, 97.1 % purity) was obtained as a white solid.

#### Synthesis of fluorescent dimers of NKp46 Bicycle peptides and NK-TICA molecule

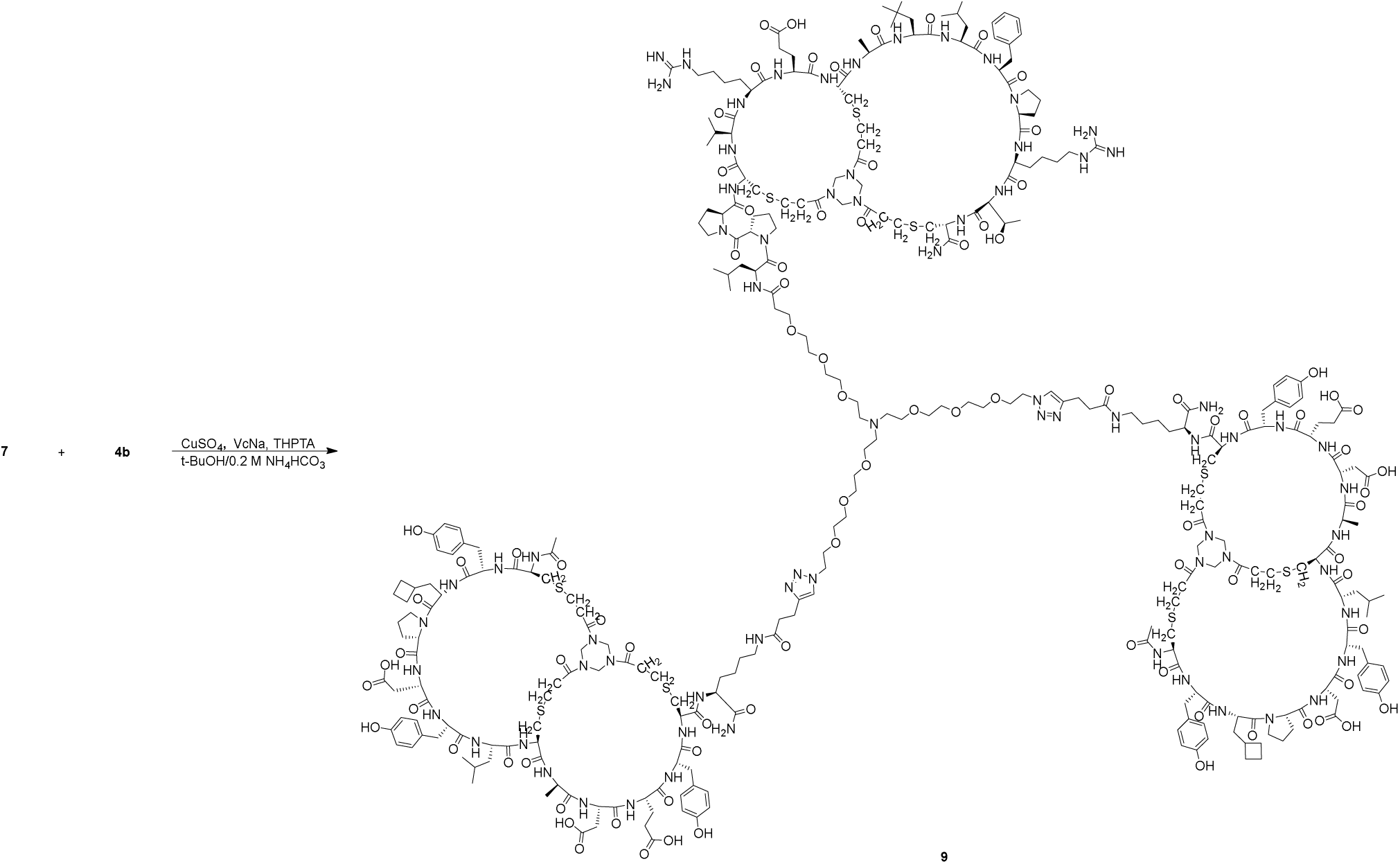

A mixture of compound 10 (5.00 mg, 9.08 µmol, 1.0 eq.), Bicycle monomer, 4b, (39.6 mg, 19.1 µmol, 2.1 eq.), and THPTA (7.90 mg, 18.2 µmol, 2.0 eq.) was dissolved in t-BuOH/H_2_O (1:1, 0.5 mL, pre-degassed and purged with N_2_ for 3 times), and then an aqueous solution of CuSO_4_ (0.4 M, 22.7µL, 1.0 eq.) and sodium ascorbate (VcNa) (3.60 mg, 18.2 µmol, 2.0 eq.) were added under N_2_. The pH of this solution was adjusted to 8 by dropwise addition of 0.2 M NH_4_HCO_3_ (in 1:1 t-BuOH/H_2_O), and the solution turned to light yellow. The reaction was stirred at 25-30 °C for 1 hr under N_2_ atmosphere. LC-MS showed compound 10 was consumed completely, and one main peak with desired m/z (calculated MW: 4713.33, observed m/z: 1178.9 ([M+4H]^4+^)) was detected. The reaction mixture was filtered to remove the undissolved residue. The crude was purified by prep-HPLC, and compound 11 (20.0 mg, 4.24 µmol, 46.7 % yield, 92.3 % purity) was obtained as a white solid.

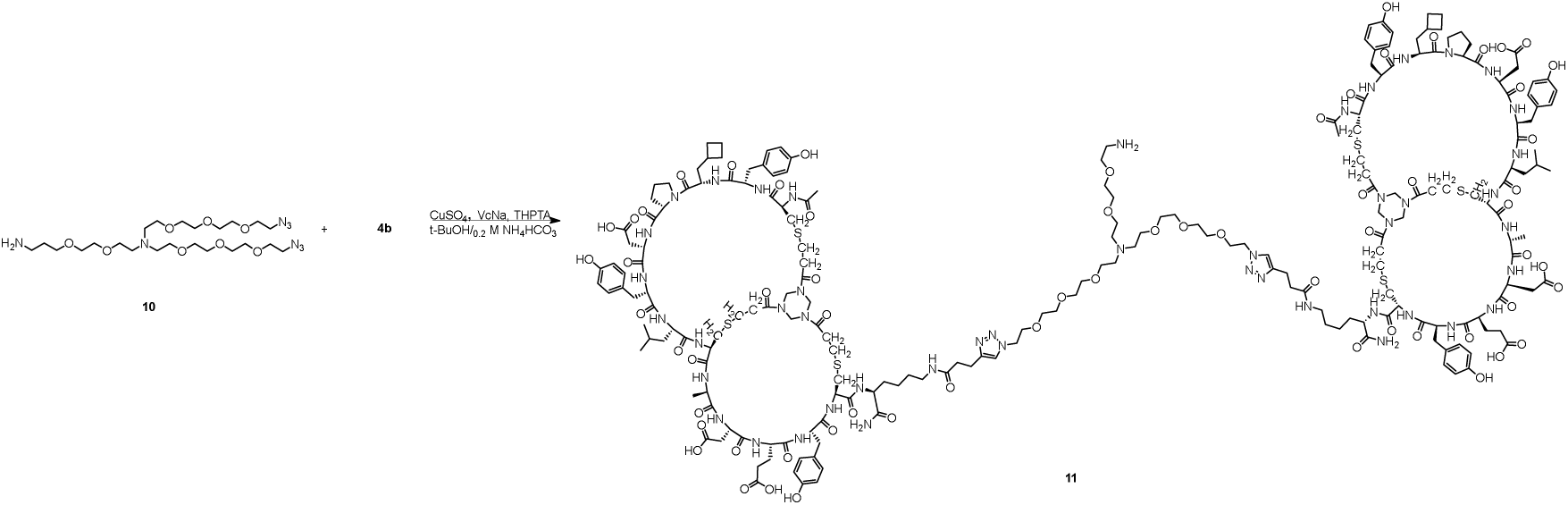

A mixture of compound 11 (20.0 mg, 4.24 µmol, 1.0 eq.), compound 12 (6.20 mg, 6.36 µmol, 1.5 eq.) and DIEA (2.20 µL, 12.7 µmol, 3.0 eq.) was dissolved in DMF (0.5 mL). The reaction stirred at 25-30 °C for 0.5 hr. LC-MS showed compound 11 was consumed completely and one main peak with desired m/z (MW: 5568.3, observed m/z: 1393.0 ([M+4H]^4+^)) was detected. The reaction mixture was filtered to remove the undissolved residue. The crude was then purified by prep-HPLC to give dye-conjugated Bicycle dimer 13 (14.2 mg, 2.40 µmol, 56.5 % yield, 94.0% purity) as a blue solid.

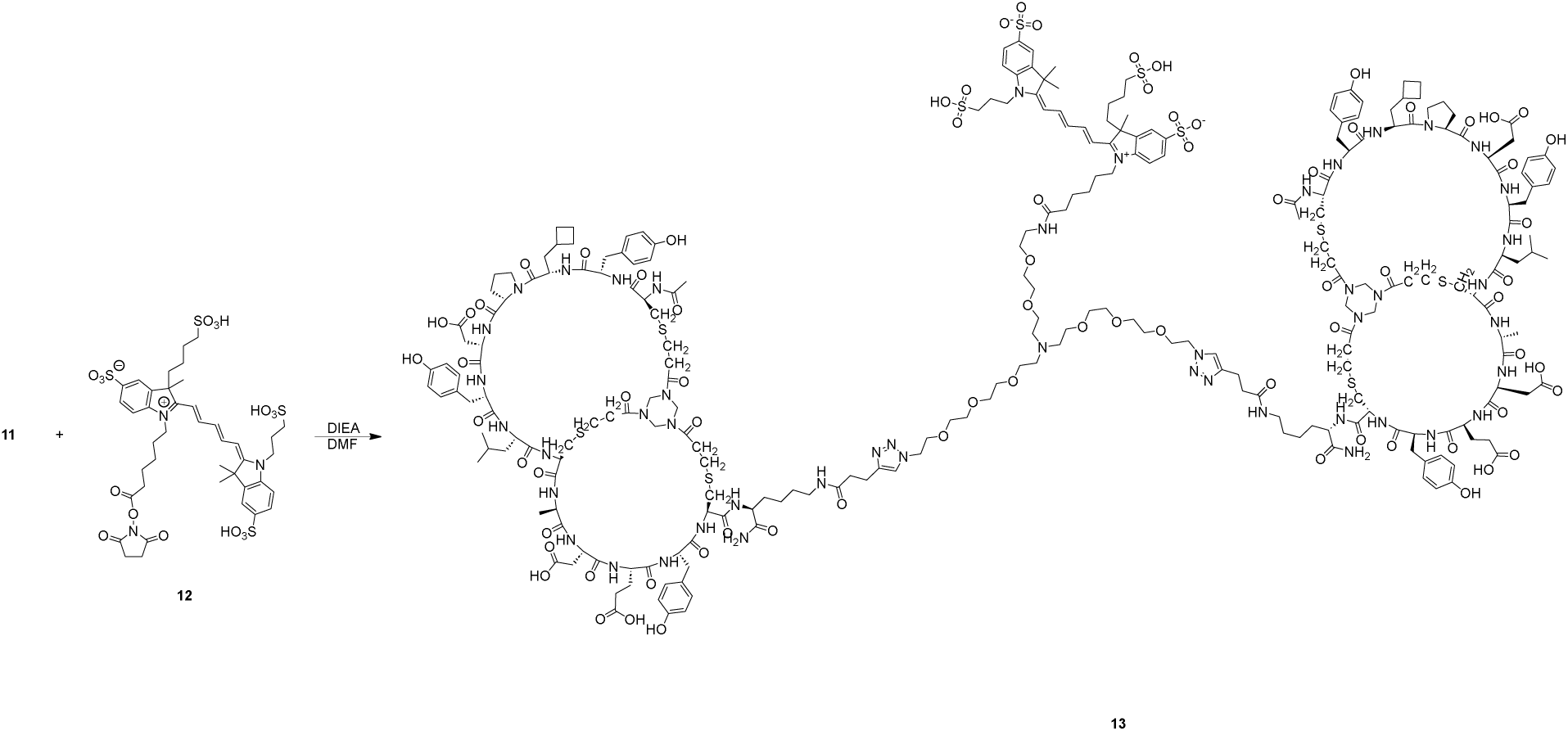

To a solution of 14 (15.0 mg, 48.4 µmol, 1.00 eq) in DMF (0.30 mL) was added DIEA (25.3 µL, 145 µmol, 3.00 eq), then 15 (54.7 mg, 58.1 µmol, 1.20 eq) was added and the mixture was stirred at 20 °C for 1 hr. LC-MS showed the PEG 14 was consumed completely and one main peak with desired m/z (MW: 1136.33, observed m/z: 1136.5 ([M+H]^+^)) was detected. The reaction mixture was purified by prep-HPLC (TFA conditions: A: 0.075 % TFA in H_2_O, B: ACN) to afford the dye conjugate 16 (40.0 mg) as a blue solid. This material was used in the following step.

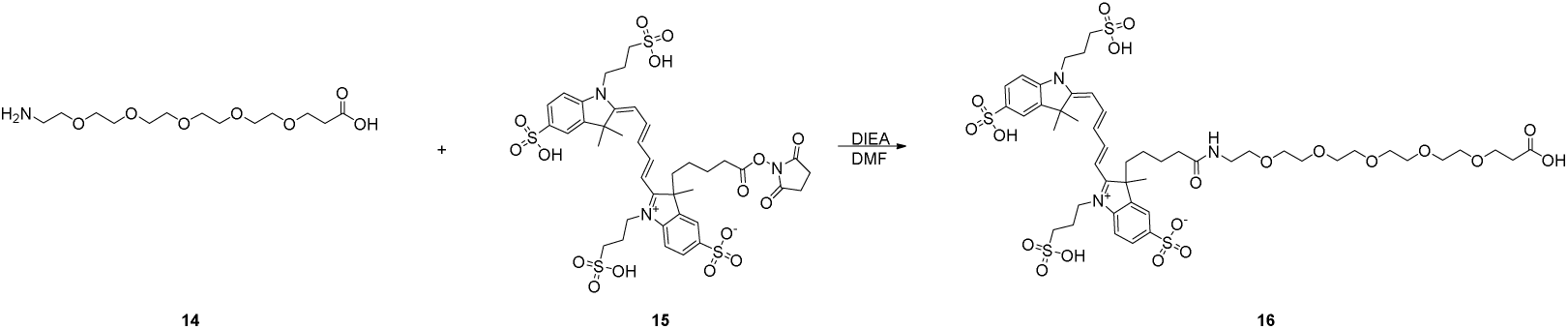

To a solution of 16 (17.2 mg, 15.1 µmol, 1.50 eq) in DMF (0.30 mL) was added DIEA (5.27 µL, 30.2 µmol, 3.00 eq) and HATU (4.22 mg, 11.1µmol, 1.10 eq) and stirred 6.00 min, then 17 (23.7 mg, 11.1 µmol, 1.10 eq) was added and the mixture was stirred at 20 °C for 24 min. LC-MS showed 16 was consumed completely and one main peak with desired m/z (MW: 3256.75, observed m/z: 1629.3([M+2H]^2+^)) was detected. The reaction mixture was purified by prep-HPLC (TFA conditions: A: 0.075 % TFA in H_2_O, B: ACN) to afford compound 18 (12.1 mg) was obtained as a blue solid.

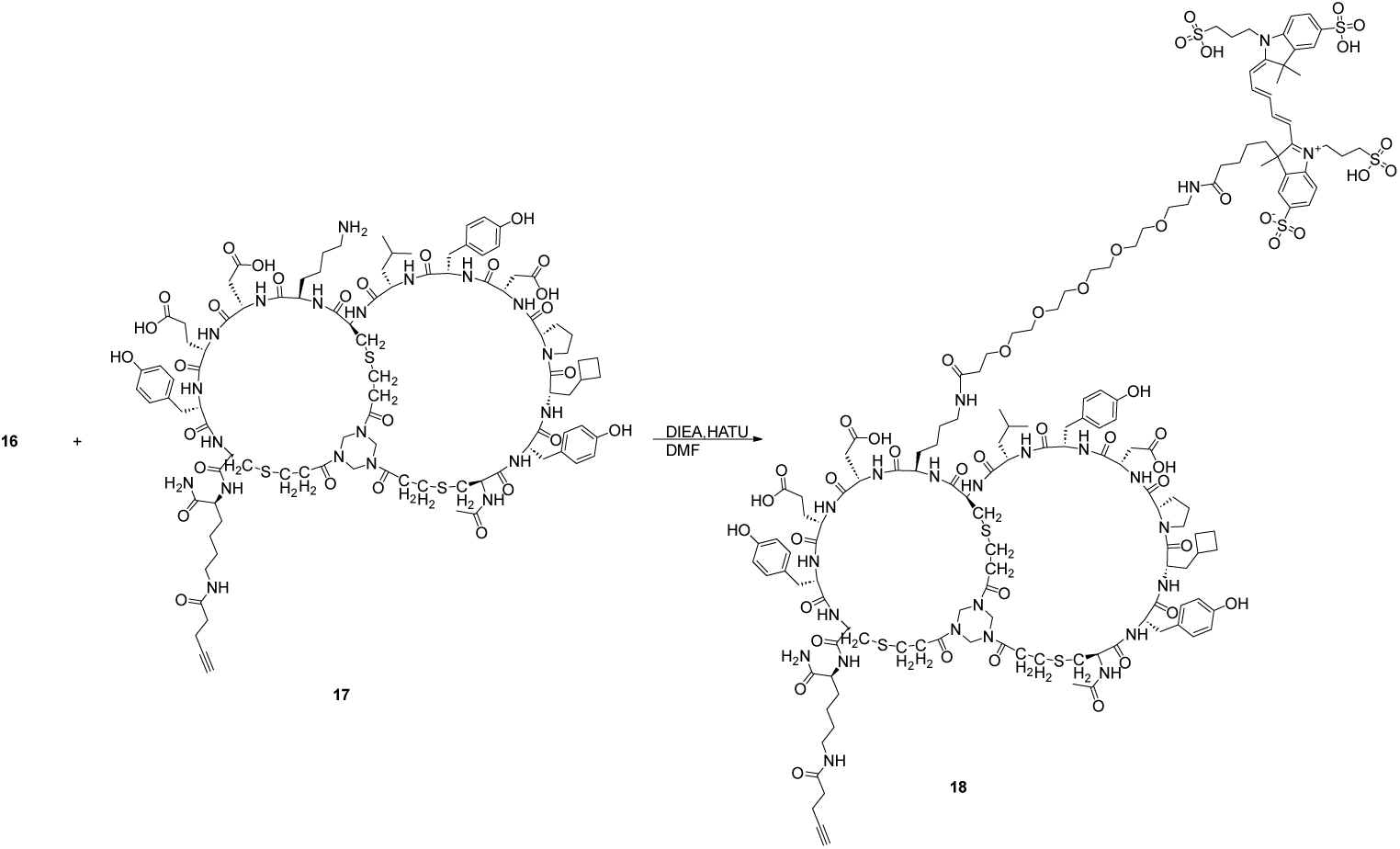

A mixture of PEG-EphA2 Bicycle conjugate peptide 3, (60.0 mg, 19.9 µmol, 1.00 eq), NKp46 Bicycle peptide 4b (45.6 mg, 21.9 µmol, 1.10 eq) and THPTA (8.67 mg, 19.9 µmol, 1.00 eq) was dissolved in t-BuOH (0.15 mL) and H_2_O (0.15 mL) (pre-degassed and purged with N_2_ for 3 times). Then aqueous solution of CuSO_4_ (0.40 M, 49.8 µL, 1.00 eq) and sodium ascorbate (VcNa) (7.91 mg, 39.9 µmol, 2.00 eq) were added under N_2_. The pH of this solution was adjusted to 8 by dropwise addition of 0.20 M NH_4_HCO_3_ (in 1:1 t-BuOH/H_2_O), and the solution turned to light yellow. The reaction mixture was stirred at 20 °C for 1 hr under N_2_ atmosphere. LC-MS showed conjugate 3 was consumed completely and one main peak with desired m/z (MW: 5087.82, observed m/z: 1697.1 ([M+3H]^3+^), 1273.1 ([M+4H]^4+^)) was detected. The reaction mixture was purified by prep-HPLC (TFA condition: A: 0.075 % TFA in H_2_O, B: ACN) to afford the conjugated heterodimer 19 (24.9 mg) as a white solid. This material was used in the next step.

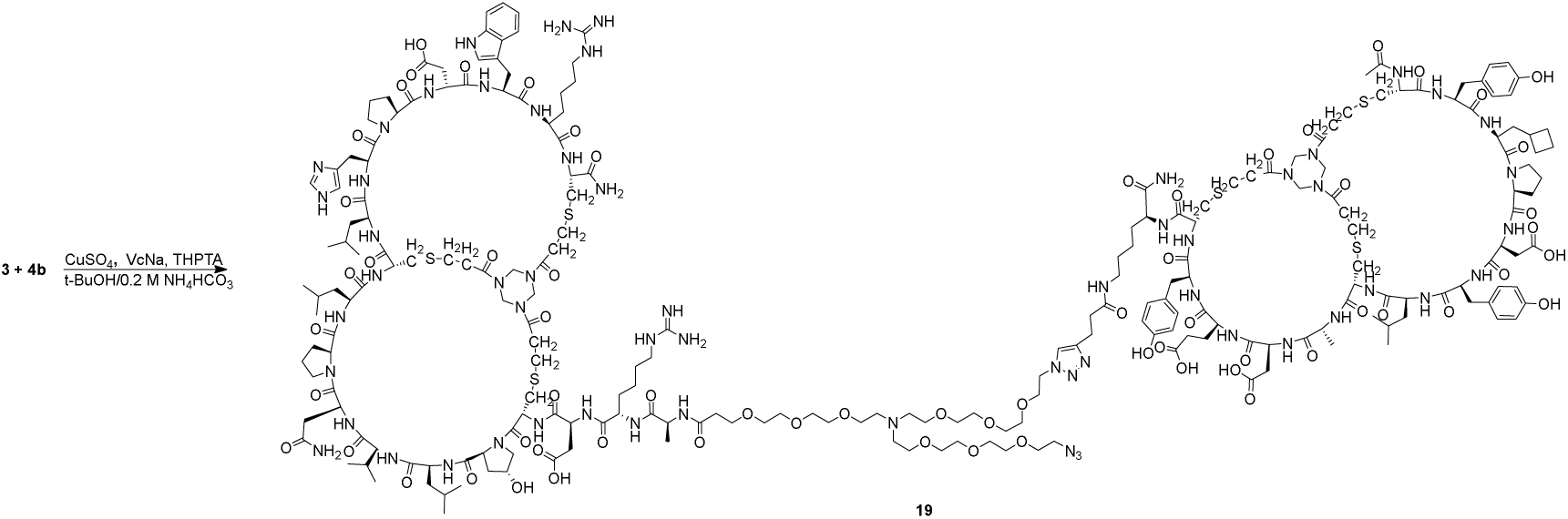

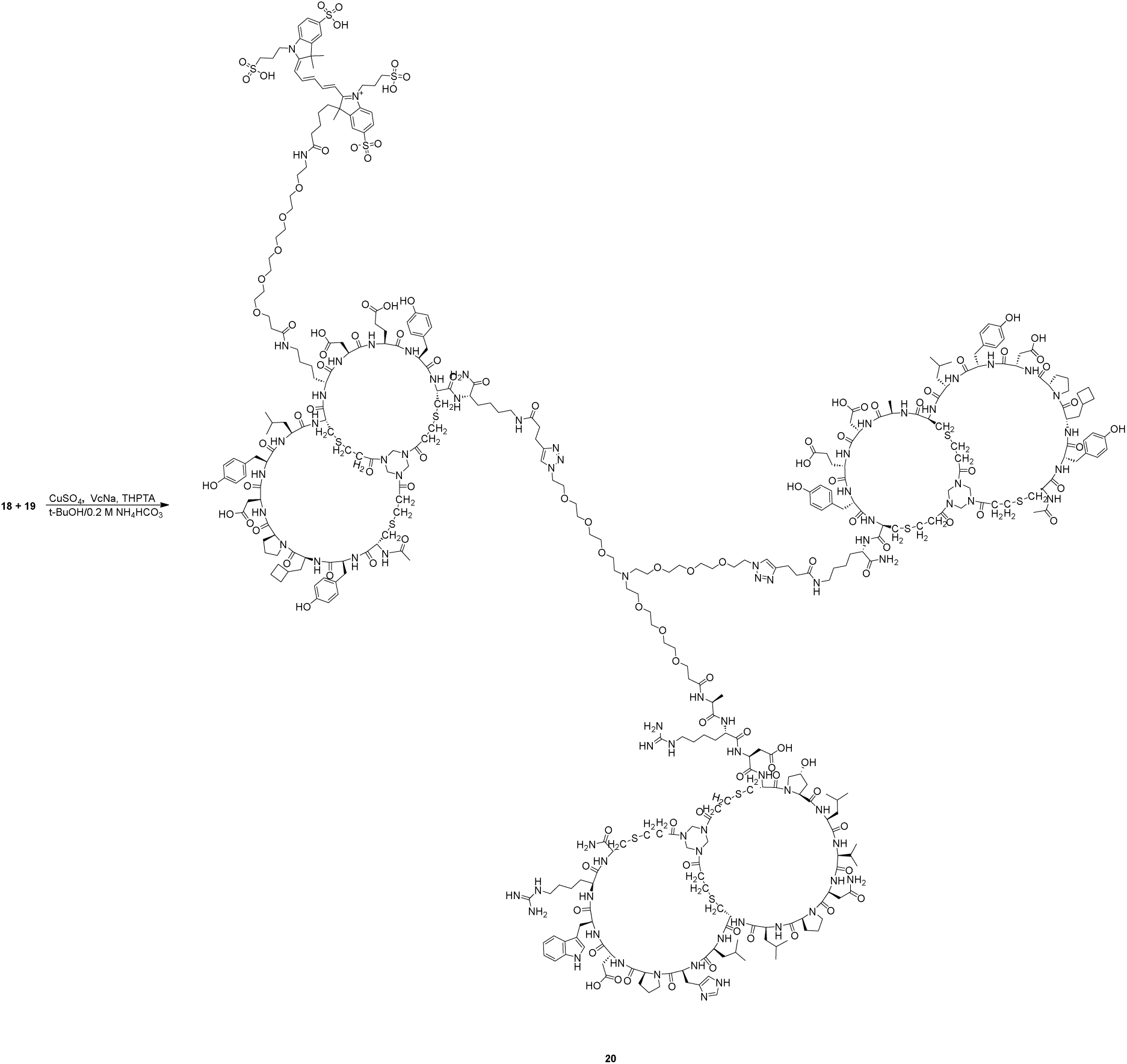

A mixture of 18 (20.7 mg, 4.09 µmol, 1.10 eq), 19 (12.1 mg, 3.72 µmol, 1.00 eq) and THPTA (1.61 mg, 3.72 µmol, 1.00 eq) was dissolved in t-BuOH (0.15 mL) and H_2_O (0.15 mL) (pre-degassed and purged with N_2_ for 3 times). Then aqueous solution of CuSO_4_ (0.40 M, 9.29 µL, 1.00 eq) and sodium ascorbate (VcNa) (1.47 mg, 7.43 µmol, 2.00 eq) were added under N_2_. The pH of this solution was adjusted to 8 by dropwise addition of 0.20 M NH_4_HCO_3_ (in 1:1 t-BuOH/H_2_O). The reaction mixture was stirred at 20 °C for 1 hr under N_2_ atmosphere. LC-MS showed compound 19 was consumed completely and one main peak with desired m/z (MW: 8344.56, observed m/z: 1669.8 ([M+5H]^5+^), 1391.9 ([M+6H]^6+^), 1193.0 ([M+7H]^7+^)) was detected. The reaction mixture was filtered to remove the undissolved residue. The crude product was purified by prep-HPLC (TFA condition: A: 0.075 % TFA in H_2_O, B: ACN) to afford 20 (15.6 mg, 1.86 µmol, 49.9 % yield, 99.3 % purity) as a blue solid.

**Figure S1.**
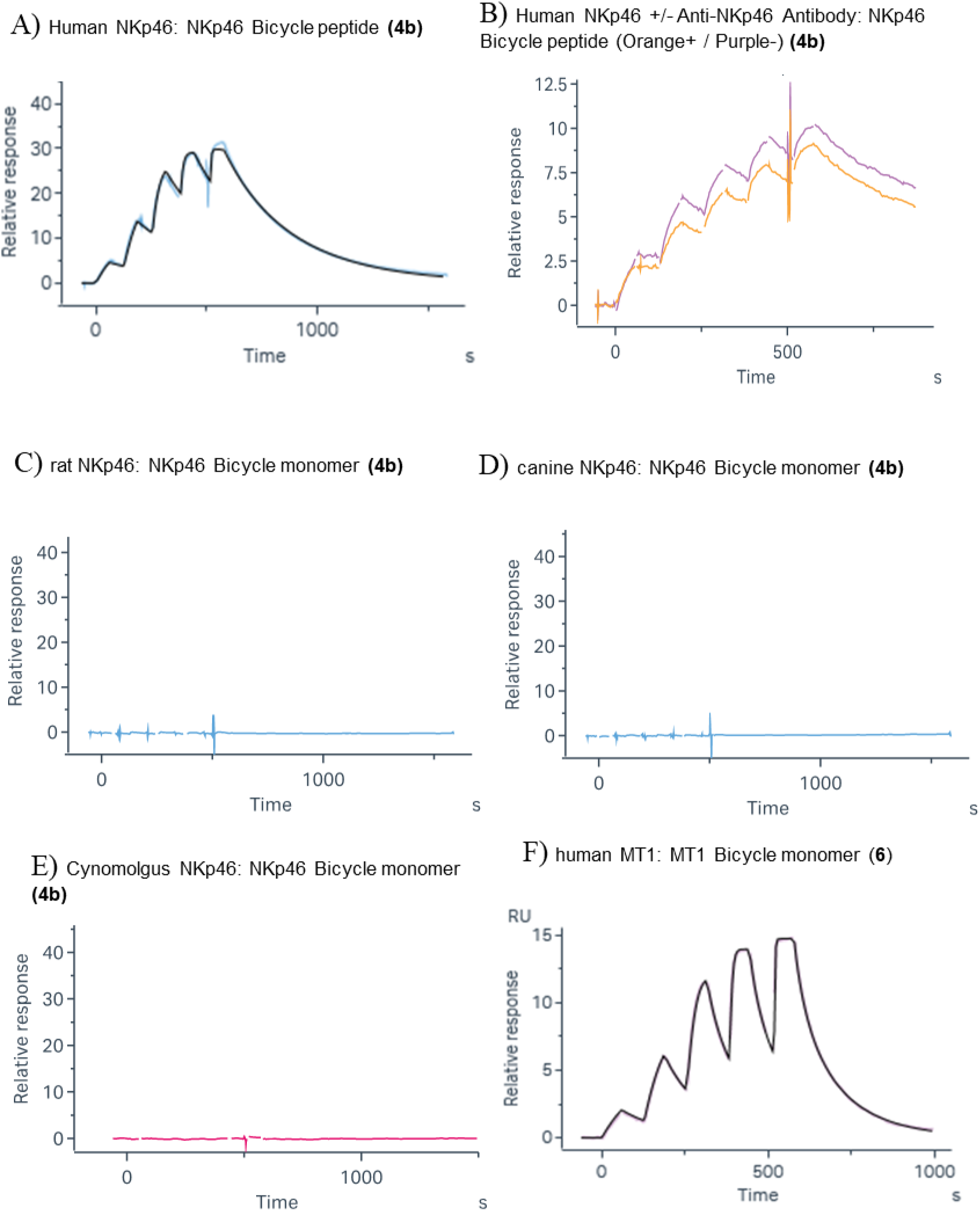

## Notes

### Summary of Updates

In the first version uploaded, two authors (SU and WHZ) were omitted in error. This has been corrected in the present version. There are no other changes to the manuscript.

